# The vast majority of somatic mutations in plants are layer-specific

**DOI:** 10.1101/2024.01.04.573414

**Authors:** Manish Goel, José A. Campoy, Kristin Krause, Lisa C. Baus, Anshupa Sahu, Hequan Sun, Birgit Walkemeier, Magdalena Marek, Randy Beaudry, David Ruiz, Bruno Huettel, Korbinian Schneeberger

## Abstract

**Background:** All plant tissues and organs develop from meristems. Plant meristems are structured organs consisting of distinct layers of stem cells. Somatic mutations occurring in one of these layers can propagate into large sectors of the plant. However, the frequency and characteristics of meristematic mutations that form the basis of somaclonal phenotypic variation remain unclear.

**Results:** Here, we analysed the frequency and distribution of somatic mutations in an individual Apricot tree. For this, we sequenced the genomes of fruit samples corresponding to distinct meristematic cell layers selected across the entire tree. Most somatic mutations (>90%) were specific to individual layers. Genotyping the somatic mutations in leaves sampled next to the fruits confirmed their meristematic origin. Interestingly, layer 1 (epidermis) had a higher mutation load than layer 2 (mesocarp), implying differential mutational dynamics between the layers. The somatic mutations followed the branching pattern of the tree. These factors led to the unexpected observation that the layer 1 samples from different branches were more similar to each other than to layer 2 samples of the same branch. Further, using single-cell RNA sequencing, we demonstrated that the layer-specific mutant alleles could only be found in the transcripts of the respective, layer-specific cell clusters and could form the basis for somaclonal phenotypic variation.

**Conclusions:** Here, we analyzed the prevalence and distribution of somatic mutations with meristematic origin. Our insights into the yet unexplored layer-specificity of such somatic mutations outlined how they can be identified and how they impact the breeding of clonally propagated crops.

## Introduction

Plant meristems are specialized tissues maintaining undifferentiated stem cells. They facilitate growth by continuously dividing and producing new cells that undergo differentiation leading to the formation of new branches and organs. Meristems are broadly structured in different zones known as tunica and corpus (1–3). Typically, the tunica consists of two peripheral layers (L1 and L2) while the corpus consists of one interior layer (L3) (4,5). During the development of new organs, the identities of the different layers are mostly conserved implying that newly formed organs are also formed of three, mostly distinct cell lineages or layers (6,7). For example, in leaves, the epidermis is formed of cells that developed from L1, mesophyll cells are from L2, and cells of the vascular tissue are from L3.

While somatic mutations in differentiated tissue will only affect small parts of a tree, somatic mutations in stem cells can be propagated to large sectors of the plant leading to genomic mosaicism between entire branches or organs (3,8). Such meristematic mutations can induce bud sport mutants, where significant parts of the plant appear or behave differently than the rest of the tree, including traits with agronomical value. Consequently, bud sports (and the underlying somatic mutations) are frequently used to improve crop species, including fruit trees, where conventional breeding based on the introgression of additional traits is slow and tedious (9–13).

Multiple studies have analysed genomic mosaicism within individual plants and discussed various aspects of somatic mutations like mutation rates, allelic frequencies, genome-wide distribution, and distribution across branches (14–22). However, most analyses were limited to bulked samples where cells from all layers were sequenced and analysed together. This can miss certain layer-specific mutations and limits our understanding of the prevalence and spectra of somatic mutations in specific cell layers (13).

Here, we exemplify the genomic heterogeneity between cell layers caused by somatic mutations in the meristem by analyzing an apricot tree. We identified and characterised somatic mutations within fruits and leaves sampled from the tips of multiple branches. Analysing layer-enriched tissues revealed an unexpectedly high load of layer-specific mutations, which are usually hidden in typical somatic mutation analyses based on bulked samples. We found that samples from the same layers were more similar than samples from the same branch. We confirmed the meristematic origin and layer-specific identity of mutations by analysing the distribution of individual mutations between the neighboring organs and using single-cell transcriptomics analysis of the leaves. Our results provide a holistic understanding of the genomic heterogeneity between the different meristematic cell lineages that remain the building blocks of plants’ genomic architecture.

## Results and Discussion

### Identifying somatic mutations in a fruit tree

*Prunus armeniaca* cultivar (cv.) Rojo Pasión is a registered apricot variety that originated from a cross between *P. armeniaca* cv. Orange Red and cv. Currot in 1996 (23). The original Rojo Pasión tree was clonally propagated for the first time in 2001. The clonal progeny were re-propagated again in 2009. For this study, we sampled from a Rojo Pasión tree of the second clonal generation growing in an orchard near Murcia, Spain. This tree is of particular interest as some of its branches show unusually low chill requirements and an early-flowering phenotype (24). Its primary stem branched into three secondary stems, which further divided into several branches. In the spring of 2020, we collected leaf and fruit samples from the tips of seven branches (Fig. 1A) (Methods). We first generated a chromosome-level haplotype resolved genome assembly of the diploid genome of Rojo Pasión using DNA from leaves of several branches. PacBio HiFi reads were separated into two sets of reads derived from either the maternal or paternal haplotypes using the trio-binning approach (25,26). The read sets were independently assembled into a haplotype-resolved, chromosome-level assembly (*k-mer*-based phasing accuracy: 99.99%, assembly completeness: >98%, assembly QV: >45) (Additional file 1: Supplementary Methods, Supplementary Figures S1-3) (27,28). The two haplotype-specific assemblies were 233.2 and 234.8 Mbp long (estimated genome size: ∼242.5 Mbp) with NG50 values of 26.9 and 28.0 Mbp for the haplotypes derived from Currot and Orange Red, respectively. We annotated 28,355 and 28,473 genes in the two genomes corresponding to a BUSCO completeness score of 96.8% for both haplotypes (Additional file 1: Supplementary Methods) (29).

**Fig. 1.**
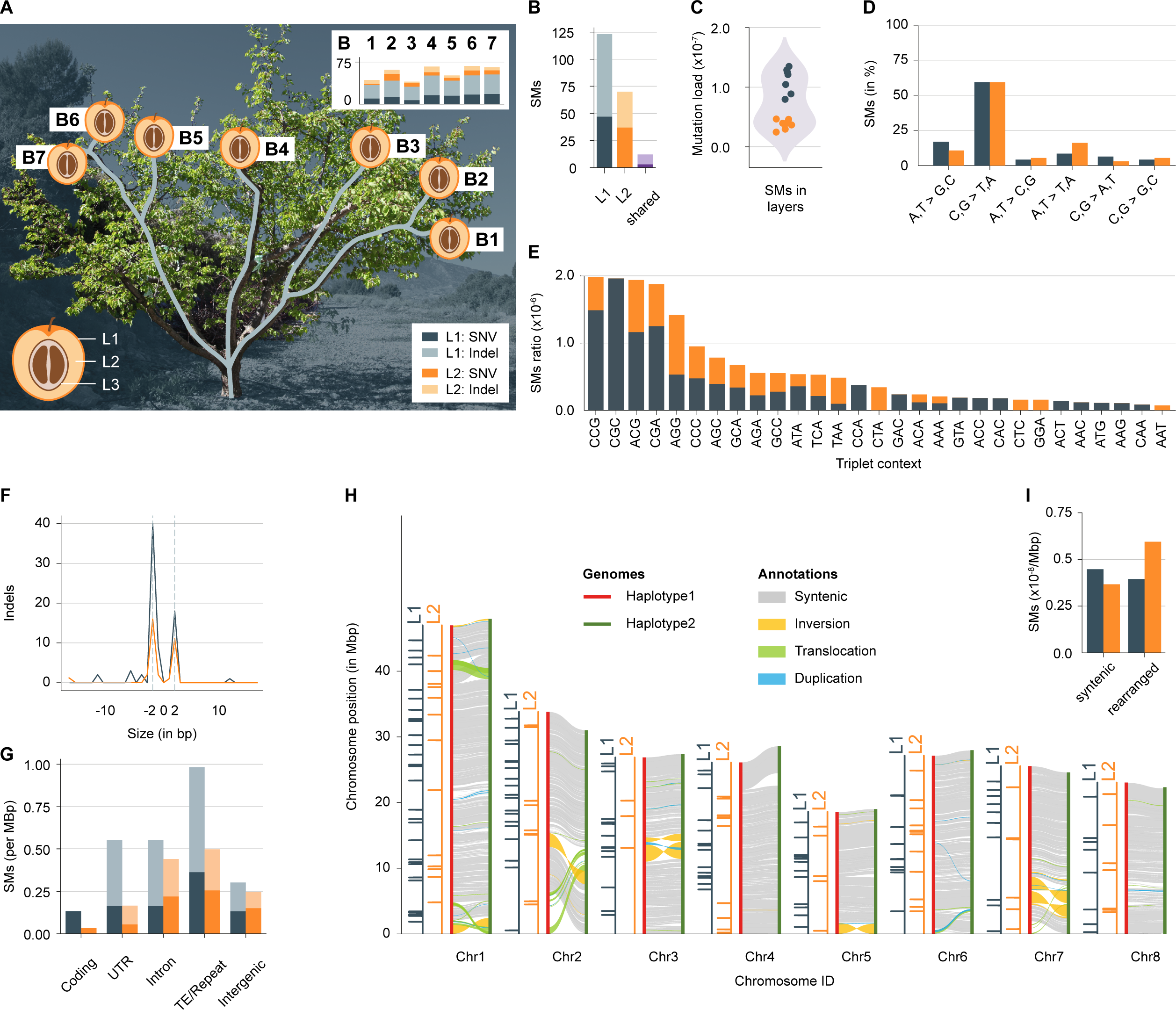
Layer-specific somatic mutations in an Apricot tree. **A)** Schematic layout of the seven branches in the apricot tree. The apricot fruit peel consists of L1, mesocarp of L2, and pre-endocarp of L3 cells. Bar plots show the number of somatic mutations identified in each branch with mutations in L1 (grey) and L2 (orange). SNVs have a dark hue while indels are light. **B)** Total number of somatic mutations (SMs) in L1, L2 or shared. **C)** Mutation load in L1 and L2. The Y-axis corresponds to the number of mutations per base in the haploid genome. **D)** Percentage of transition and transversion point mutations. **E)** Triplet context of point mutations (the middle base is mutated and the triplets are shown using the canonical form). **F)** Indel size distribution. Negative values correspond to deletion whereas positive values correspond to insertions. **G)** Normalised somatic mutation frequency in different genomic regions. UTR: untranslated regions; TE: transposable element. Values on the y-axis were normalised by the total length of the genomic region where somatic mutations could be identified. **H)** Plotsr visualisation of structural differences between the two haplotypes and the distribution of somatic mutations across the chromosomes. **I)** Normalised somatic mutation frequency in syntenic and structurally rearranged regions. Values on the y-axis were normalised by the total number of mutations identified in the layer.

We dissected the peel and the flesh of individual fruits from seven branches to generate layer-enriched samples. The peel samples corresponded to cells from L1 and the flesh samples to cells from L2 (Fig. 1A, Methods) (7). In apricot fruits, cells from L3 are restricted to a thin layer of the endocarp making it challenging to extract them without contamination from other layers. Therefore, we did not consider L3 in this study. We also collected seven individual leaves adjacent to each of the selected fruits. The DNA of these 21 samples (14 from fruits and seven from leaves) were sequenced using Illumina paired-end sequencing with very high sequence coverage ranging from 198 to 422x per sample (Methods, Additional file 1: Supplementary Methods, Additional file 2: Supplementary Table S1).

After aligning the sequencing reads to the new assembly, we combined multiple pipelines to identify somatic mutations in the fruits (Methods, Additional file 1: Supplementary Figures S4-5, Additional file 2: Supplementary Table S2-S6). Overall, we identified 215 small *de novo* mutations (point mutations and small indels), six loss of heterozygosity (LOH) mutations (where heterozygous alleles changed into homozygous alleles), and two complex mutations (Additional file 2: Supplementary Table S3-6). Despite thorough screening, no transposable element movement or meiosis-like chromosome arm exchanges could be identified (14). To confirm that the somatic mutations were real, we targeted the validation of 20 somatic mutations with digital PCR. Of those, we could confirm 14 mutations, while for the remaining six either the primers or probes were inconclusive (Additional file 1: Supplementary Methods, Supplementary Figures S6-S8, Additional file 2: Supplementary Table S7-S10).

### Most of the somatic mutations are specific to individual layers

Among the 215 small-scale mutations (point mutations and small indels), 205 were found in the layer-enriched samples of the fruits while the remaining were found in leaves only (discussed in the following sections).

The vast majority of these 205 mutations in fruits were specific to individual layers (94% (n=193)), while only 6% (n=12) were shared between L1 and L2 samples. Likewise, all six LOH mutations were exclusively found in L1 samples. The low number of shared mutations suggested low but existing cellular exchange between the meristematic cell layers (3,30). However, we cannot exclude that the shared mutations resulted from imperfections in the generation of the layer-enriched samples, in particular as half of the shared mutations were observed at a single branch suggesting that the separation of the cell lineages at this branch was not perfect.

Among the layer-specific mutations, there were significantly more mutations (64% (n=123)) in L1 as compared to L2 (36% (n=70)) (Fig. 1B). The higher mutation load in L1 was consistent across all branches suggesting different mutational processes in the layers of the meristem and pointing to a relaxed control of genome integrity in L1 (Fig. 1A, C) (31).

The 193 layer-specific mutations in L1 or L2 included 84 point mutations and 109 small indels. The point mutations were enriched for transitions with a strong bias for C to T and G to A mutations as described for mutations in plants before (Fig. 1D) (14,16,32,33). Interestingly, however, this bias was not only found in the mutations of L2 (which can be propagated to the next generation during sexual reproduction) but also among the mutations in L1 (which are not propagated to the next generation during sexual reproduction) (6). Most of these mutations occurred at CpG sites, the most common DNA methylation context in plants, which can trigger the formation of C to T mutations (Fig. 1E) (33,34). In plants, UV-B based DNA damage is associated with mutations, however, the somatic mutations, including those specific to the outer L1 layer, were not enriched for mutation types associated with UV-B (35). Like the point mutations, indel mutations also showed a strong bias in their spectrum. Around 70% (76 of the 109) of the indels were 2bp long and occurred in AT dinucleotide microsatellite regions (Fig. 1F). These mutations were potentially introduced by DNA polymerase slippage (36,37).

Most somatic mutations were found in transposable elements and repeats (Additional file 1: Supplementary Figure S9) (14). When normalized for genomic abundance and callability of genomic regions, we found that in L1 transposable elements had a significantly higher mutation rate (right tail Fisher’s exact test, p-adjusted for L1: 0.0005) while coding regions featured a significantly lower mutation rate in both L1 and L2 as compared to a random distribution of mutations (left tail Fisher’s exact test, p-adjusted for L1: 0.006, L2: 0.008) (Fig. 1G). This is consistent with a recent report suggesting lower mutation rates in evolutionarily conserved regions (38), however, in some parts, the under-representation of mutations in genes could also be explained by an under-representation of microsatellite regions within them (37).

We also checked the effect of local sequence diversity between the haplotypes (heterozygosity) on the frequency of somatic mutations as it has been reported that local heterozygosity could lead to regionally increased mutation rates (39). For both, L1 and L2 however, somatic mutations were distributed without any recognizable effect of the large-scale or small-scale nucleotide diversity (Fig. 1H, I, Additional file 1: Supplementary Figure S10) (40,41).

### The distribution of somatic mutations throughout the tree

The formation of new branches starts with the development of new axillary meristems from the founding cells of the apical meristem (42). If the founding cells carry somatic mutations, the newly formed axillary meristem will inherit and propagate them into the new branch (17).

To understand the propagation of mutations within the tree, we analysed the distribution of the somatic mutations across different branches (Fig. 2A) (13,43). We found that 31% (64 out of 205) of the somatic mutations were present in multiple branches. Almost all of the shared mutations (94% (n=60)) were found in neighbouring branches where their sharing patterns between the branches agreed with the tree topology (Additional file 1: Supplementary Figure S11). Only four mutations did not follow this pattern. Of these, two were found in B2 and B3 but not in B1, one was found in B1 and B4 but not in B2 and B3, and the remaining mutation was found in all branches except for one. While the distribution of the somatic mutations strongly agreed with the tree topology, branch length did not correlate with the number of somatic mutations (Additional file 1: Supplementary Figure S12).

**Fig. 2.**
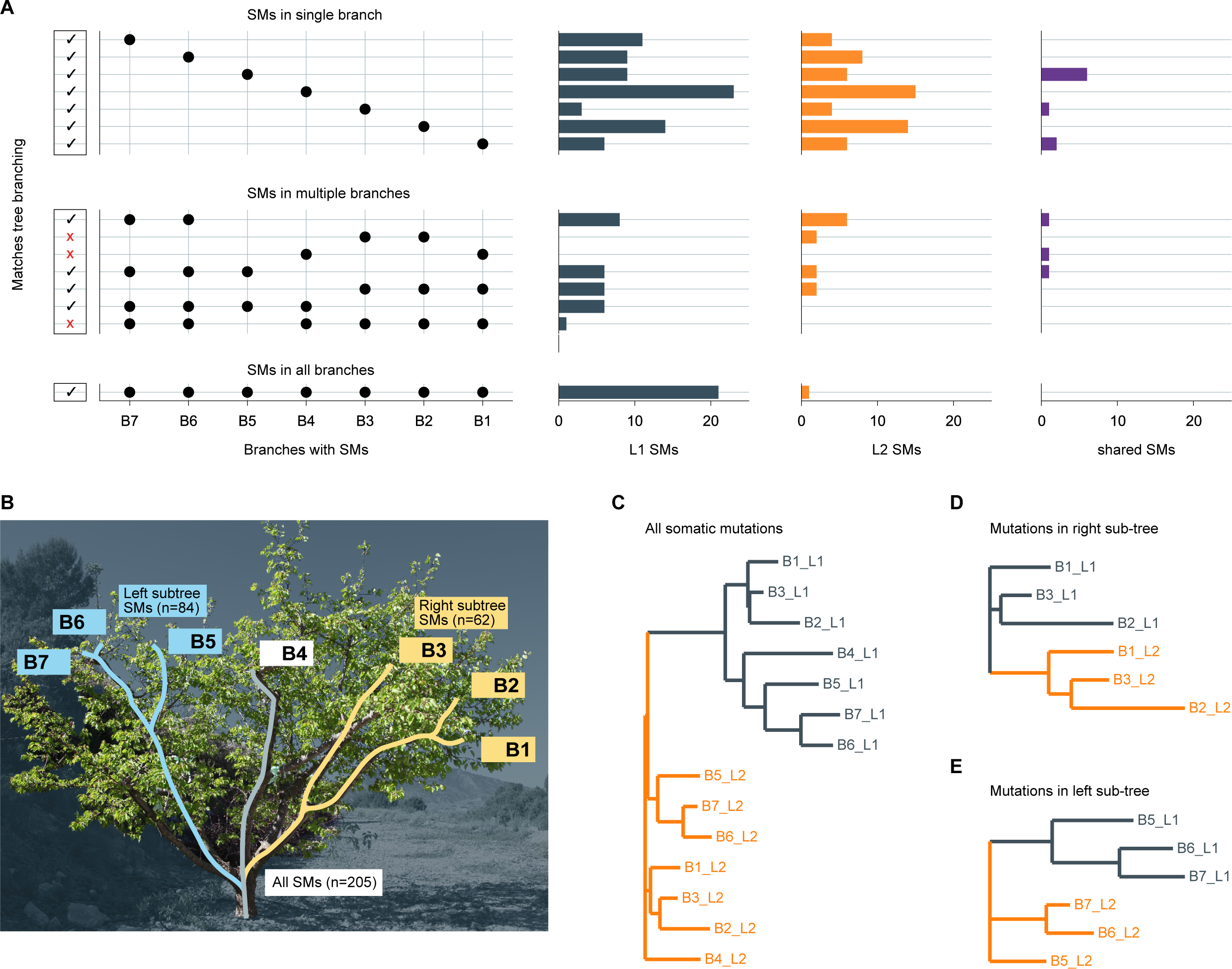
Somatic mutations sharing between layers and branches. **A)** The distribution of somatic mutations across branches. The five panels show (left to right) i) whether the mutations follow the topology of the tree (black checkmark) or not (red cross), ii) the sets of branches in which mutations are present, each row corresponds to a set and the black point marks the branches present in the set, iii-v) the number of somatic mutations in L1, L2, or shared between layers. **B)** Somatic mutations in the entire tree and the two sub-trees. **C-E)** Unrooted Neighbouring Joining trees of the individual samples outlining the similarity between the samples. **C)** All samples. **D)** Samples only from the sub-tree including branches B1-B3. **E)** Samples only from the sub-tree including branches B5-B7.

The somatic mutations in L1 and L2 were similarly distributed across the tree, with the remarkable exception that 21 mutations in L1 were present in all seven branches while only one such mutation was found in L2 (Fig. 2A, Additional file 1: Supplementary Figure S13). The high number of mutations occurring in all branches (compared to mutations in six, five, or four branches) suggested that they might be clonally inherited from the mother tree and did not occur during the development of this tree. Further, the presence of these mutations in all branches also suggested that they were already fixed in the cutting that was used for the grafting event from which this focal tree grew. The observation that more L1 mutations are shared across all branches is in agreement with a higher mutation load in L1 as this implies that more L1 mutations are carried over clonal generations as well.

Unexpectedly, samples of the same layers were more similar to each other as compared to the samples of the same branch (but from different layers) This implied that samples from L1 of different branches were more similar to each other than to L2 samples of the same tissue, and vice versa (Fig. 2B, C). To exclude that this pattern was solely introduced by the somatic mutations that were shared across all branches, we separately analysed only those mutations that originated in this tree (by focusing on two sub-trees B1-B3 and B5-B7) (Fig. 2D, E, Additional file 1: Supplementary Figure S14). Again, the L1 samples were more related to each other than the L1 and L2 samples of the same branch, and vice versa.

Taken together, our results suggest that mutations in all layers are propagated during branching and clonal reproduction. Unlike sexual reproduction, where only mutations in L2 can be inherited by the next generation, clonal reproduction can propagate mutations in all layers. Additionally, mutations, that are fixed in individual branches used for clonal reproduction, will get fixed in the entire tree within the next generation. This will result in the accumulation of mutations within the plants of the next generation.

### Mutation sharing between neighbouring organs reveals their meristematic origin

In addition to the 64 somatic mutations that were shared between different branches, we also found 141 mutations specific to individual branches. The high number of branch-specific mutations contrasts the high probability of mutations growing into newly formed branches, questioning whether these mutations truly occurred in the meristems or during the development of these fruits.

To understand the origin of these somatic mutations in more detail, we sequenced the DNA of seven leaves that grew adjacent to the sequenced fruits. While leaf tissue also includes all three cell layers, we did not generate layer-specific samples, instead, we deeply sequenced the DNA from bulked leaf cells (sequence coverage from 198-422x). To note, bulked leaf samples do not include equal amounts of cells of the different layers, but each layer is present with different amounts of cells (44).

We identified 70 somatic mutations within the seven leaves (Fig. 3A) (Methods, Additional file 2: Supplementary Table S3). Most of these mutations were found in the adjacent fruits (85% (n=60)). The remaining mutations (15% (n=10)) were specific to leaves (Fig. 3B). The leaf-specific mutations were also absent in all other samples of the tree suggesting that they originated in the individual leaves.

**Fig. 3.**
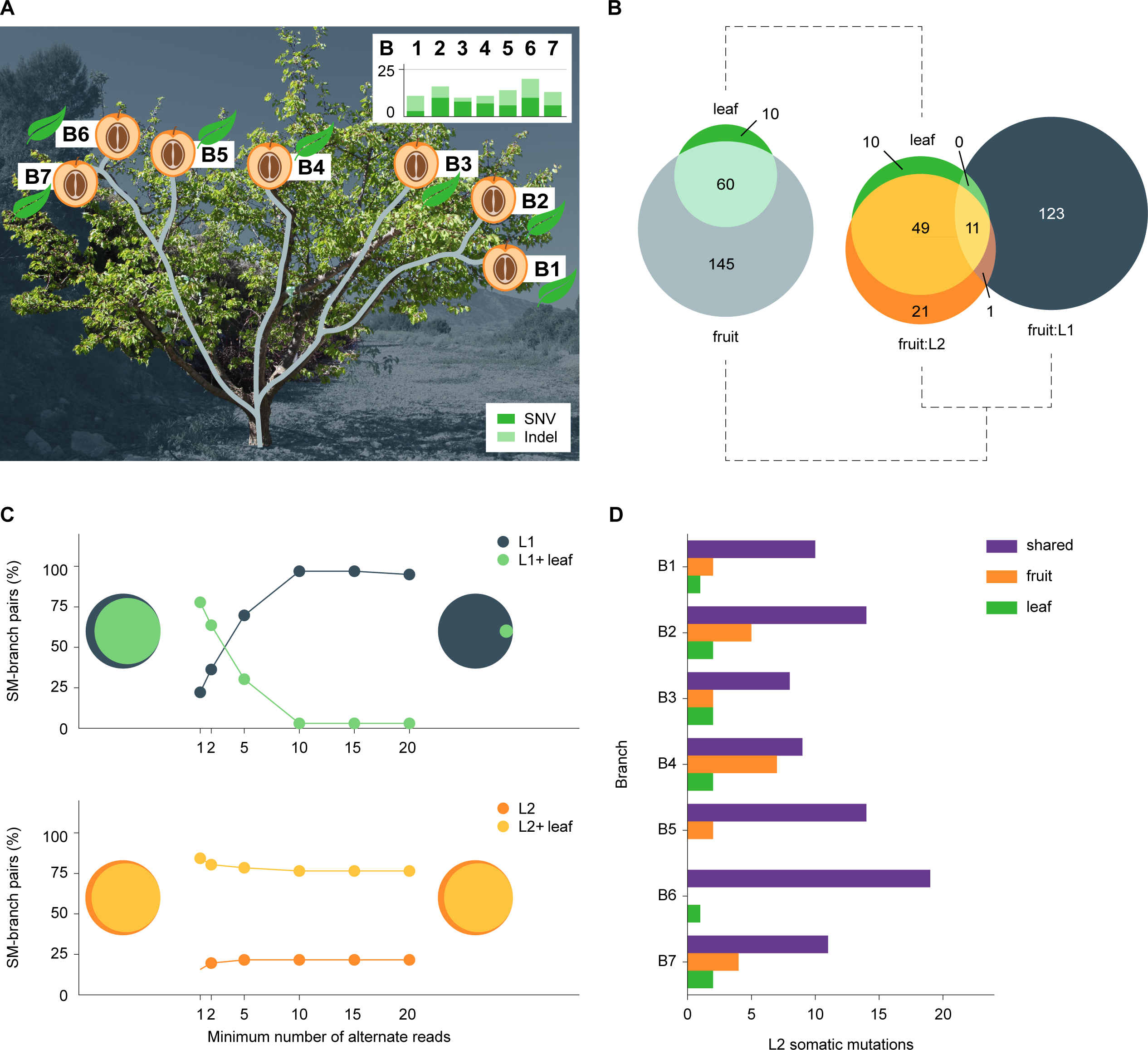
Genomic differences between adjacent fruit and leaf pairs. **A)** Somatic mutations were identified in leaves adjacent to the selected fruits. The bar plots show the number of somatic mutations identified in each leaf with SNVs in dark green and indels in light green. **B)** Both Venn diagrams show the number of somatic mutations shared between fruits and leaves. Left: all mutations; right: somatic mutations in the fruits are shown separately for L1 and L2. **C)** Technical evaluation of the impact of different read-support requirements on genotyping somatic mutations. Upper panel: successful genotyping of 50 known somatic mutations (n=50) in the L1 of the fruits (grey) as well as in the L1 of the fruits and the leaf samples in parallel (only if both samples support it, we considered the mutations being positively genotyped) (lightgreen). For each somatic mutation, only branches in which the somatic mutations were originally identified in, were used for the genotyping. The X-axis shows the minimal-read-support required to find support for the genotype a somatic mutation. The Y-axis shows the percentage of tests in which somatic mutations were positively genotyped in only L1 or both (L1 and leaf). The Venn diagrams on the left and right show the sharing of somatic mutations between L1 and leaf for read cut-offs of 1 and 20, respectively. Lower panel: similar plot for genotyping somatic mutations (n=40) identified in L2. **D)** Number of putative L2 somatic mutations (X-axis) present in fruit, leaf, or both.

In contrast, only 30% (60 out of 205) of the mutations in fruits were shared with the neighbouring leaves including a striking difference between the individual layers. While 70% (49 out of 70) of the L2-specific mutations in the fruits were found in leaves, not a single L1-specific mutation in the fruits was identified in leaves (0 out of 123) (Fig. 3B).

This difference in mutation sharing between the different layers could result from an under-representation of L1 cells in the leaf samples (Additional file 1: Supplementary Figure S15). To test this, we performed a targeted search for the mutations found in L1 in the fruits (Additional file 1: Supplementary Methods). We re-analysed the sequencing data of the leaves by lowering the read support required to genotype a layer-specific mutation and found weak evidence for ∼80% of the L1 mutations (Fig. 3C). Lowering the read support requirement during *de novo* mutation identification (to one or two reads used for genotyping) would have led to an extremely high false positive rate, which in turn would make it impossible to find the set of true mutations (45,46). On the contrary, the number of mutations, that could be found in the L2 of the fruits and their corresponding leaves, was mostly unaffected by changes in read support cut-off. This implied that almost all L2 mutations shared between leaves and fruits were already identified (Fig. 3C).

Although we found evidence for 80% of the L1 mutations in the leaves, it is possible that some of them resulted from false positives or sequencing errors. Therefore, to understand how many mutations were truly shared between closely neighbouring organs, we zoomed in on L2 mutations as those were most reliably identified in both organs. To validate the presence/absence of the L2 mutations in both sample types, we compared the read frequencies of the mutations in the fruit and leaf samples. The frequencies of the mutations that were shared in both organs were very similar (Additional file 1: Supplementary Figure S16, S17), however, mutations that were specific to either L2 layers or leaves had almost no read support in the respective other samples. This supported the annotation of the presence and absence of the mutations.

Overall, we found that 65% (60 out of 92) of the mutations identified in L2 in fruits or in the leaves were shared between the neighbouring organs and were of putative meristematic origin. This ratio was mostly conserved across all seven branches (Fig. 3D). The remaining mutations (35% (n=32)) were observed in either the leaves or the fruits only. All these 32 mutations were restricted to individual branches and were not found in other branches. This suggests that these mutations either happened during the development of the specific organs or that the mutations were absent in the founder cells from which the individual neighboring organs developed (43).

Taken together, by comparing mutations in neighboring organs, we found that at least two-thirds of the somatic mutations were shared between the organs, implying their meristematic origin. For the remaining mutations, we could not distinguish if they occurred in the meristems or the organs.

### Meristematic mutations affect transcripts only in the respective cell layer

Layer-specific, meristematic mutations can affect the function of cells that originated from respective layers (47). Mutations in layer 2, for example, are expected to affect all the different cell types within the mesophyll of a leaf, while all other cells should not be affected.

To test if meristematic mutations affect only cell types of the mutated layer, we sequenced the transcriptomes of the same leaves that we used earlier for mutation identification. Conventional bulk RNA sequencing of entire leaves, however, combines the transcripts of different cell populations including the cells from different layers. Therefore, to identify and analyse the transcription of specific somatic mutations in individual cell populations, we used single-cell RNA sequencing for four different branches (B2, B4, B5, and B6) (Fig. 4A, Methods, Additional file 1: Supplementary Methods).

**Fig. 4.**
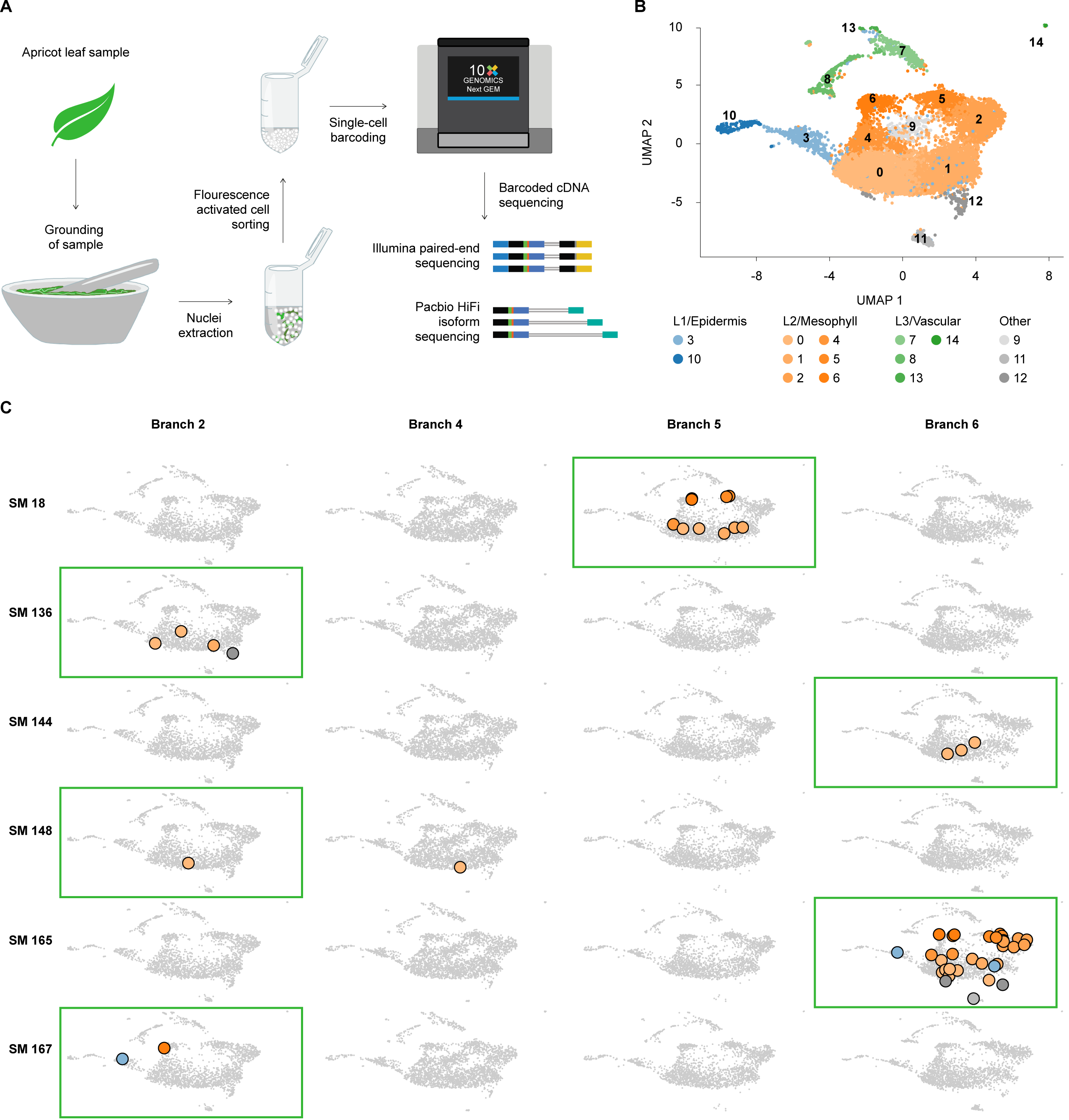
Layer-specific transcription of somatic mutations. **A)** Workflow for generating scRNA-seq and scISO-seq data from single leaves. **B)** Expression atlas of apricot leaves. The 15 clusters are numbered 0-14. Clusters from L1, L2, and L3 are coloured with blue, orange, and green shades, respectively. Clusters corresponding to micro-environmental and developmental variations, which were mostly specific to individual branches, are coloured in shades of grey. **C)** Transcription of mutated alleles. The figure highlights cells with transcripts carrying mutant alleles for six somatic mutations. The four columns correspond to four branches. Each row corresponds to one somatic mutation. Green rectangles highlight the branches in which the corresponding somatic mutations were identified and where the mutant alleles are expected. Grey cells do not have any reads with the mutant allele, whereas cells with mutant alleles are coloured based on the expression cluster they were assigned to in B.

The four datasets consisted of a total of 10,293 cells corresponding to fifteen different transcription clusters (Fig. 4B). The clusters were classified as epidermis (L1 consisting of two transcription clusters), mesophyll (L2; six clusters) or vascular (L3; four clusters) based on cell-type specific marker genes, gene set enrichment analysis and comparison to other leaf expression atlases (Methods, Additional file 2: Supplementary Table S11-S15) (48–50). Most cells corresponded to mesophyll (78.2% (n=8051)), while the epidermis (8.9% (n=917)) and vascular (7.2% (n=744)) were represented by fewer cells (Additional file 1: Supplementary Figure S18). The low representation of L1 cells (epidermis) within the leaves’ samples corroborated our earlier observation that it is difficult to find somatic mutations in the L1 layer in bulked leaf tissue.

The remaining cells (5.6% (n=581)) were found in three small clusters specific to individual branches. One of these clusters consisted of cells expressing biotic stress response genes while the other clusters showed gene expression patterns similar to dividing cells. These branch-specific clusters reflect the differences in the micro-environment and developmental stages of the different leaves and outline the sampling biases in single-cell experiments from wild collections. The scRNA-seq libraries, which are usually sequenced with short reads, typically reveal only small parts of a transcript, making searching for somatic mutations inefficient (51). To analyse entire transcripts, we also sequenced each library with PacBio HiFi long-read sequencing (scISO-seq) (Fig. 4A, Methods, Additional file 1: Supplementary Methods). This approach ensured that the transcripts from the same cells had the same barcodes in both the scRNA-seq and the scISO-seq datasets, thus allowing for expression-based clustering (generated using the scRNA-seq datasets) and to search for mutations based on full-length sequenced transcripts (using the scISO-seq datasets) (Additional file 1: Supplementary Figure S19). The additional scISO-seq allowed analysis of 8.2 to 10.5 Mb (12.5–20%) additional genome for the presence of somatic mutations, as compared to analyzing the scRNA-seq alone (Additional file 1: Supplementary Figure S20).

Overall, we identified mutant alleles for six somatic point mutations in these four branches (Fig. 4C). All six mutations were originally identified in L2. Intriguingly, all mutant alleles were almost exclusively found in transcripts of cells assigned to L2 clusters in the branches in which they were originally identified (Fig. 4C). Overall, 48 L2 cells featured mutant alleles, while only three L1 and none of the L3 cells featured reads with mutant alleles.

Two of the mutations were highly expressed (SM18 and SM165 in Fig. 4C). Cells carrying one of these two mutant alleles could be found in most clusters assigned to L2 (SM18 was found in five and SM165 was found in all six clusters of L2). This suggests that all six transcription clusters (cell types) were truly derived from L2.

In contrast, the four other mutations that were less strongly expressed, were only found in a few cells. This was due to the overall low expression and not due to genetic heterogeneity in the L2 cells as the number of cells with mutant alleles matched the number of cells with the wild-type alleles (Additional file 1: Supplementary Figure S21, S22).

The separation of expression clusters and their assignment to the individual cell layers allowed us to analyze expression differences between the individual layers. As somatic mutations often result from inefficient DNA repair, it is possible that the high mutation load in L1 as compared to L2 resulted from differences in DNA repair between the layers (31). When checking for gene expression differences of the Apricot orthologs of *A. thaliana* DNA repair genes between the layers in our scRNA-seq dataset, we however, did not find significant differences (Additional file 1: Supplementary Figure S23).

In summary, with the help of single-cell expression data, we confirmed the layer-specificity of the somatic mutations and increased the resolution of analysing the distribution of somatic mutations within complex tissues. We found meristematic mutations (once propagated into developed tissue) retain their layer-specific identity and occur in all the cell lineages developed from the respective meristematic layers.

## Conclusions

Here we have analysed the frequency, distribution, and transcription of somatic mutations in different cell layers of an apricot tree. The different layers have their origin in the architecture of the meristem. The meristem is organised in layers leading to distinct cell lineages in the newly formed branches and organs. In each layer, we found distinct sets of somatic mutations with only a small proportion of mutations shared between them. Intriguingly, the samples of the same layer were more similar to each other as compared to samples of the same branch. This suggested that the individual layers of the tree grew almost independently and that selection on somatic cells would act independently on the different layers. This effect would be stronger for the L1 as cellular migration between L2 and L3 could result in multiple low-allele frequency mutations but fewer fixed mutations (52,53). Our observations, however, contrast some earlier reports where the distribution of the somatic mutations did not follow the branching topology of the tree (22). Our study included a few differences from these studies though. For example, we excluded very low allele frequency mutations. Low allele frequency mutations are less likely to be of meristemic origin and are prone to systematic errors during sequencing and variant calling. This can lead to a discrepancy between the topology of the tree and the observed mutations. In future, it will thus be interesting to study the outgrowth probability of mutations out of the meristem (*e.g.*, introduced through spatial biases in the distribution of mutations within the meristem).

Layer 1 (epidermis) of the fruit tree revealed significantly more mutations than layer 2. This is similar to a recent report in potato, where regenerated plants revealed different mutational loads depending on the tissue they were regenerated from (54). This also aligns well with the frequent identification of bud sports specific to layer 1 in various fruit trees (13). However, as the identification of mutations in layer 1 is difficult (as outlined here), the genetic basis of only a few sport mutants has been uncovered until today.

In general, clonal reproduction will lead to an accumulation of layer-specific mutations resulting in increased genomic divergence between different tissues. Moreover, in the absence of meiotic recombination, this will include the accumulation of deleterious mutations which could be the basis for the frequently reported “collapse” of specific cultivars after several rounds of clonal reproduction. Altogether this suggests that the analysis of individual cell layers is advantageous to understand the presence and distribution of somatic mutations. Although less straightforward than analysing bulked samples, layer-specific analyses can reveal the previously hidden spectrum of mutations, offering a holistic perspective on their accumulation and propagation.

## Methods

### Plant material for mutation analysis

We collected single leaves and fruits from the tip of seven different branches of a diploid Apricot tree of the *Rojo Pasión* variety that originated from a cross between the Orange Red and Currot ecotypes and grows in an orchard close to Murcia, Spain. The samples were collected into Falcon tubes and directly snap-frozen in liquid nitrogen. Distances between sampling points were measured using a rope that was scaled in centimetres.

### Apricot layer-specific dissection from fruits and sequencing

Fruits were 1-3 cm in diameter and frozen at -80°C. One fruit from the tip of each branch was used (except B6 for which two adjacent fruits were used because there was not sufficient plant material available in one fruit). For the dissection, they were kept in liquid nitrogen. The fruits were held by fingers wearing nitrile gloves over cotton wool gloves. The surface was scratched with a fresh small scalpel to get layer 1. The material was put immediately in a 2ml Eppendorf tube, which was sitting in a styrofoam rack over liquid nitrogen to snap freeze. The fruit was peeled again (1-2 mm thick) to remove any leftover layer 1 cells and the suture region was discarded. Then the fruit was put back into the liquid nitrogen as quickly as possible. The fruit was cut into quarters. Three of these quarters were frozen again. Next, the mesocarp (green central part of the fruit) was dissected and put into 2ml or 5ml Eppendorf tubes, according to size, to get layer 2.

Layer 1 was ground with a steel bead in a Retsch Mill, followed by grinding with a small pistil. Layer 2 was ground with liquid nitrogen in a mortar and weighed for approximately 100mg in one 2ml Eppendorf tube. The DNA was extracted with Machery Nagel Kit NucleoSpin PlantII. Elution with 40µl PE elution buffer. After the first elution, it was eluted again with the eluate. For the quality of the DNA, the OD260/280 and OD260/280 were measured with NanoDrop. For the exact concentration, the DNA was measured with Qubit.

The extracted DNA samples were sent to Novogene for library preparation and sequencing using Illumina paired-end sequencing platforms.

### Leaf genome and transcriptome sequencing

For four branches (B2, B4, B5, B6), we used a single leaf each to perform single-cell whole-genome sequencing (scDNA-seq), single-cell RNA-sequencing (scRNA-seq), and single-cell Isoform sequencing (scISO-seq). Sequencing reads from scDNA-seq sequencing were used in bulk and individual cell-level analysis was not considered. For the other three branches (B1, B3, B7), single leaves were used for conventional whole-genome sequencing. The corresponding methods are described in Additional file 1: Supplementary Methods.

### Somatic mutation identification using layer-enriched fruit DNA

Raw reads were trimmed using skewer (-r 0.1 -d 0.05 -k 8 -q 20 -l 75 -m -pe) and aligned to the genome assembly of the Currot haplotype using bowtie2 with --end-to-end and --very-sensitive preset (55,56). Duplicated reads were marked and removed using samtools (57). Allelic read counts were identified at all positions using bam-readcount (-b 30 -q 10) (58). This distribution of read counts after filtering was used to select the optimal read depth range for each sample (Additional file 1: Supplementary Figure S4). Positions with at least three reads with a non-reference allele were selected using in-house scripts as candidate somatic mutation loci.

Somatic mutations were then identified using custom pipelines. The first pipeline identified layer-specific mutations in individual branches. This pipeline filtered out positions that were 1) heterozygous between the two haplotypes (identified using syri (40)), 2) outside optimal read depth range (Additional file 2: Supplementary Table S2), 3) heterozygous in both L1 and L2 of a branch (alternate allele frequency (AF) between 0.3 and 0.65), 4) had small AF difference between layers (AF difference < 0.25), 5) consisted of noisy alignments. Positions supported by 20 alternate reads and had high AF difference (>99th percentile of AF difference between L1 and L2 in the branch) were selected as candidate somatic mutations and were manually curated using IGV (59). The second pipeline identified layer-specific somatic mutations that were shared between branches. This pipeline filtered out positions that were 1) heterozygous between the two haplotypes, 2) outside optimal read depth range, 3) present in only one branch, 4) supported by few reads with alternate alleles (read count <= 20) in all samples, 5) having low log^2^ fold-change between the mean alternate AF in the layers. The remaining positions were manually curated using IGV to select somatic mutations. The third pipeline identified layer-specific somatic mutations in a branch using fold-change ratios between alternate allele frequencies. This pipeline filtered out positions that were 1) heterozygous between the two haplotypes, 2) outside optimal read depth range, 3) present in more than one branch, 4) supported by few alternate reads (read count <= 20), 5) had low log^2^ fold-change between layers. Selected candidates were manually curated using IGV. Somatic mutations identified by these pipelines were combined to get the list of layer-specific mutations. Next, we identified somatic mutations that were shared between layers. For this, we detected mutations shared between the 14 samples without considering the sample layer as a factor. However, analysing all samples together meant that we could not easily filter out the background noise in sequencing data, therefore we had to modify the pipeline to remove putative false-positive mutations. This pipeline filters out positions that were 1) heterozygous between the two haplotypes, 2) not supported by at least 20 reads with alternate alleles or had alternate allele frequency < 0.25 in all samples, 3) supported by the alternate allele in all samples, 4) repetitive (read-depth > 500), 5) supported by alternate allele reads only from a single chromosome strand. The selected positions were manually curated using IGV.

### Identifying loss of heterozygosity mutations

We checked SNVs and indels between the haplotypes for loss of heterozygosity (LOH) mutations. Two signals were used: allele frequency differences between layers and reads with switching haplotypes. In each branch, we first identified the number of reads aligned at the short variant sites and calculated the alternate allele frequency. We filtered out positions that 1) were heterozygous in both L1 and L2 (0.35 <= AF <= 0.6), 2) had low AF in both L1 and L2 (AF <= 0.1), 3) had absolute allele frequency difference between L1 and L2 of less than 0.25, 4) were sequenced with few reads (shallow read-depth). The remaining positions were candidate LOH mutations based on allele frequency differences. Next, for each sample, pileup data was analysed and positions where at least 10 reads had haplotype switch (Currot to Orange Red, or vice-versa) between short variant positions, that were within 1kb of each other, were selected. This provided a clear signal of LOH and allowed the removal of false positives. Positions with haplotype switches in both L1 and L2 were filtered out. Positions showing both allele frequency differences and haplotype switching were manually curated using IGV.

### Somatic mutation identification in leaves

#### Pre-processing and alignment of single-cell whole genome sequencing datasets

Raw fastq files were filtered to remove reads smaller than 50bp as cellranger-dna could not process them. The remaining reads were analysed using cellranger-dna and clustered into cells based on 10X barcodes. Cells with less than 10000 reads were filtered out and readsets corresponding to individual cells were created. Each read set was aligned to the Currot genome using bowtie2 (end-to-end alignment with –very-sensitive preset). Duplicated reads were marked and removed using samtools. Alignment BAM files from individual cells were merged using samtools to get an alignment file corresponding to the sample.

#### Pre-processing and alignment of conventional WGS datasets

Raw reads were trimmed using skewer (-r 0.1 -d 0.05 -k 8 -q 20 -l 75 -m -pe) and then aligned to the Currot genome using bowtie2 (end-to-end alignment with –very-sensitive preset). Duplicated reads were marked and removed using samtools.

#### Somatic mutation calling

Bam-readcount was used to get allelic read counts (-b 30 -q 10) and genomic positions with less than 3 reads with alternate alleles were filtered out using custom Python scripts (58). The pipeline for somatic mutation identification then filtered out positions that were 1) heterozygous between the two haplotypes, 2) outside optimal read depth range (Additional file 2: Supplementary Table S2), 3) supported by <20 alternate allele reads or <0.25 alternate allele frequency. The pipeline also filtered positions with library or sequencing biases. The remaining positions were manually curated using IGV.

The final somatic mutation list consisted of all mutations identified from the layer-enriched fruit samples and leaf samples. Further, all samples were manually re-checked at all of the identified somatic mutation positions, and if there was sufficient evidence that a somatic mutation could be present in the sample but currently not called, then such somatic mutations were added in the sample as well.

### Single-cell RNA-seq analysis

Raw reads were processed using cellranger-7.1.0 (from 10x Genomics) using the ‘count’ subcommand with ‘—expect-cells 3500 –include-introns true’ parameters and a maximum intron size of 5000. The filtered barcode matrices generated by the cellranger were analysed using R and Seurat 4.0.5 (60,61). A sample-specific threshold was used for filtering cells based on UMI count. Only cells with unique feature counts greater than 500 and UMI count greater than 100 but less than the 95th percentile of UMI count distribution were included in the analysis. Each sample was normalized and variable features were individually identified using the *NormalizeData* and *FindVariableFeatures* function. The selection method parameter was set to vst and the nFeatures parameter was set to 2000. Genes that were repeatedly variable across samples were identified using the *SelectIntegrationFeatures* function and subsequently used for integrating the samples using the *FindIntegrationAnchors* and *IntegrateData* functions. The downstream analysis used standard Seurat workflow for visualisation and clustering with *ScaleData*, *RunPCA*, *RunUMAP*, *FindNeighbors* and *FindClusters* functions. The resolution parameter for clustering was set to 0.6. Cluster annotation was done by identifying the marker genes for individual clusters using the *FindMarkers* function.

The 3-D models of apricot proteins were generated using AlphaFold (62). Orthologous *Arabidopsis thaliana* genes were identified using sequence and structural alignments with orthofinder and foldseek (63,64). These *A.thaliana* genes were used for enrichment analysis using the StringDb R package (48). Only marker genes with a p-adjusted value less than 0.05 and a log^2^-fold change value greater than 0 were considered for enrichment analysis. We also used manually curated A.*thaliana* marker genes (Additional file 2: Supplementary Table S11) as well as those reported by Kim *et al.* 2021 and Berrío *et al.* for cluster annotation (49,50).

We extracted the barcodes for cells in each cluster using custom scripts. These were used to generate cluster-specific bam files using samtools. Bam-readcount was used to genotype these bam files at somatic mutation positions. One mutation genotyped because of mis-mapped reads was filtered out.

### Single-cell ISO-seq analysis

The scISO-seq reads were processed using the methods and workflow provided by PacBio (https://github.com/PacificBiosciences/pbbioconda, https://isoseq.how/getting-started.html). This involved, the removal of cDNA primers using *lima* (parameters: --isoseq --dump-clips --dump-removed), UMI and barcode extraction using *tag* (--design T-12U-16B), removal of polyA tail with *refine* (--require-polya), barcode correction using *correct* (--B 3M-february-2018-REVERSE-COMPLEMENTED.txt.gz -M 2 --method percentile –percentile [sample_specific_value]), and deduplication of reads using *groupdedup*. For barcode correction, reference 10x barcodes provided by PacBio were used (https://downloads.pacbcloud.com/public/dataset/MAS-Seq/REF-10x_barcodes/3M-february-2018-REVERSE-COMPLEMENTED.txt.gz). The percentile cut-off values used for correction were B2: 85, B4: 85, B5: 80, and B6: 83. The processed scISO-seq reads were divided into read sets corresponding to individual cell types using the clustering (and corresponding barcodes) generated using the scRNA-seq data. The cluster-specific read sets were aligned to the genome using minimap2 and then genotyped using bam-readcount. One mutation genotyped because of mis-mapped reads was filtered out.

## Declarations

### Ethics approval and consent to participate

Not applicable

### Consent for publication

Not applicable

### Availability of data and materials

Data generated and used as part of this study will be made available on acceptance of the manuscript at EBI (https://www.ebi.ac.uk/) under Bioproject PRJEB71142. Hi-C sequencing data used for genome assembly and RNA-seq data used for genome annotation are available under Bioproject PRJEB37669 (25). All scripts used for analysis and processing are available at https://github.com/schneebergerlab/apricot_layer_mutations.

### Competing interests

The authors declare that they have no competing interests.

### Funding

This work was funded by the “Humboldt Research Fellowship for Experienced Researchers” (Alexander von Humboldt Foundation) (JAC), the Marie Skłodowska-Curie Individual Fellowship PrunMut (789673) (JAC), Deutsche Forschungsgemeinschaft (DFG, German Research Foundation) under Germany’s Excellence Strategy—EXC 2048/1–390686111 (KS) and European Research Council (ERC) grant ‘INTERACT’ (802629) (KS).

### Authors’ contributions

The project was conceptualised and designed by MG, JAC, KK, RB, KS. Data generation and wet-lab experiments were done by JAC, KK, LCB, BW, MM. Data analysis was done by MG, AS, HS. DR, BH, and KS supervised the project. MG and KS wrote the manuscript with contributions from all authors. All authors read and approved the final manuscript.

## Acknowledgements

We would like to thank Hernán López for helpful discussions, Samija Amar for help in designing illustrations and figures, and Antonio Molina for providing access to the plant material.

## List of Additional Files

| File name | File format | Title of data | Description of data |
| --- | --- | --- | --- |
| Additional file 1 | .pdf | Additional notes and figures | Additional notes describing the method and supplementary figures |
| Additional file 2 | .xlsx | Additional tables | Supplementary data and supplementary results |

## Notes

### Competing Interest Statement

The authors have declared no competing interest.

### Summary of Updates

The new version highlights the intriguing finding that the L1 samples from different branches are more similar to each other compared to the L2 samples from the same branch. This observation is discussed within the context of the propagation of somatic mutation in clonally reproducing plants as well as during branching events in a tree.

