## Additional file 1 for "The vast majority of somatic mutations in plants are layer-specific"

### Supplementary Methods

#### Leaf PacBio HiFi sequencing

Leaves from multiple branches were pooled. High molecular weight DNA was isolated from 1.5 grams of leaf material with a NucleoBond HMW DNA kit (Macherey Nagel). Quality was assessed with a FEMTOpulse device (Agilent) and quantity was measured by fluorometry Quantus (Promega). A PacBio HiFi library was then prepared according to the manual "Procedure & Checklist - Preparing HiFi SMRTbell® Libraries using SMRTbell Express Template Prep Kit 2.0" with initial DNA fragmentation by g-Tubes (Covaris) and final library size selection was done using BluePippin (Sage Science). Size distribution was again controlled by FEMTOpulse (Agilent). Size-selected libraries were sequenced on a PacBio Sequel II device with Binding Kit 2.0 and Sequel II Sequencing Kit 2.0 for 30 h.

#### Genome assembly

The PacBio HiFi sequencing of apricot leaves resulted in 613,662 reads with a total read length of ~10Gbp approximating 20x haploid coverage for the apricot genome (expected diploid genome size: ~485Mb) (1). The HiFi reads were phased using the trio-binning approach with HiCanu and linkage groups generated previously (1–3). The phasing accuracy was calculated using Meryl and KMC (4,5). In both cases, trio-binning phased the HiFi reads better. We then assembled trio-binning phased reads using HiFiasm (6). The contig level assembly generated by HiFiasm had some duplicated regions incorrectly collapsed. To overcome this, we used the unitig assemblies generated by HiFiasm. The unitig assemblies were 296.6Mbp (1117 unitigs) and 296.0Mbp (1136 unitigs) long for the Currot and Orange Red haplotypes, respectively. Next, we manually purged the unitigs. For this, we aligned the trimmed Illumina short-reads to the unitigs (‘initial assembly’) using minimap2, processed the BAM files using samtools, and then selected unitigs that were longer than 100kb or have >30x mean read-depth using custom scripts (7–9). Then, we selected all Illumina reads that did not align with the selected unitigs and mapped them to the filtered-out unitigs. Unitigs that mapped well were re-added to the list of selected unitigs, thus creating a contig level assembly. We repeated this purging step three times resulting in the contig level assembly. The purged assemblies were 237.3 Mbp (291 contigs) and 239.4 Mbp (307 contigs) long for the Currot and Orange Red haplotypes, respectively.

The assemblies were polished using the Illumina short reads from parent trees with pilon and then three times with phased HiFi reads using racon (10,11). *K-mer-based* assembly quality statistics were generated using Merqury (4). Next, we break chimeric contigs. For this, we aligned Hi-C reads generated by Campoy *et al.* (1) to the phased assemblies using BWA (12), pre-processed the read alignments using AllHiC and bamToBed (13–15), and then called chimeric positions using SALSA (16). We checked the alignment of HiFi reads at the candidate chimeric loci using samplot (17) and loci that were not supported by HiFi reads were selected for breaking. We then filtered out organellular contigs by aligning organellular sequences from previous studies (18–20). Contigs with more than 80% region mapping to organellular contigs were filtered out. Ragtag was used to group the contigs into different linkage groups (each linkage group corresponds to one chromosome) using linkage groups generated by Campoy *et al.* as the reference (21). The contigs that could not be grouped were filtered out. Contigs from individual linkage groups were scaffolded using Hi-C reads. For this, we first aligned the Hi-C reads to the post-chimeric break diploid genomes. Then the reads were split into different linkage-group read-sets based on the linkage group of the contig to which the read aligned. Reads from the same linkage group were aligned to corresponding contigs followed by the standard AllHiC scaffolding pipeline (13). AllHiC was run with k=1, to get one chromosome from the contigs of the linkage group. The scaffolds generated by AllHiC were manually curated and incorrect joins were corrected. Finally, the chromosomes were oriented and named to match the assembly from (1) and the ungrouped contigs were added back.

#### Genome annotation

We used a similar pipeline as used by Campoy *et al.* (1). In short, this pipeline integrates *ab initio* gene predictions using Augustus, GlimmerHMM, and SNAP (22–24), RNA-seq based transcript identification using StringTie (25) and protein sequence alignment-based transcript prediction using Exonerate (26). These evidences were integrated using EvidenceModeler (27). RepeatMasker was used for TE annotation as well as for filtering TE-related genes (28). The predicted gene models were improved using custom scripts and Scipio. Gene models with in-frame stop codons and CDS lengths that were not a multiple of three were removed. Gene models that were haplotype-specific but not identified in the other haplotype were added (29). Finally, PASA was run twice to add UTR regions using *de novo* transcripts from RNA-Seq reads generated by Trinity (30,31). BUSCO was used to evaluate the gene models (using the eudicots_odb10 database) and eggnog-mapper to get functional annotation of the genes (32,33).

#### Single-cell sequencing

##### Nuclei extraction from Apricot leaves

One single Apricot leaf was taken from each branch (B2, B4, B5, B6) and was ground in liquid nitrogen. The ground material was then split in half and used as input to create scDNA-seq and scRNA-seq/scISO-seq libraries, respectively, using the 10x™ Chromium controller and library-specific protocol. In brief, nuclei were extracted as follows. Ground leaf material was dissolved in 10 ml nuclear isolation buffer (NIB) (10 mM MES-KOH (pH 5.4), 10 mM NaCl, 10 mM KCl, 2.5 mM EDTA (pH 8), 250 mM sucrose, 0.1 mM spermine, 0.5 mM spermidine, 1 mM DTT, 1% PVP) and gently inverted until homogenous. Then, 2ml of TritonX-100 (10%) was added and the sample was gently inverted until homogenous and incubated on ice for 5 min. Afterwards, the sample was filtered through two layers of miracloth and centrifuged at 4°C and 800 g (breaks/acceleration at 3) for 15 min. Thereafter, the supernatant was discarded and the nuclei pellet was washed twice with 10ml NIB without TritonX-100, using the previous centrifugation settings. After the final wash, the nuclei pellet was gently dissolved in 1ml NIB and filtered through a 30 µm, 20 µm and 10 µm sieve (Sysmex CellTricks Disposable Filters), respectively, to remove debris. The filtered nuclei were then pelleted using the previous centrifugation settings and dissolved in 0.5-1 ml NIB. Afterwards, an aliquot of the nuclei sample was DAPI-stained (1-5 mg/ml) to verify nucleus integrity using a fluorescence microscope (ECLIPSE Ts2R, Nikon).

**Fluorescence-activated cell sorting**

Nuclei for scDNA-seq library preparation were DAPI-stained (1-5 mg/ml) and sorted in 0.9% NaCl sheath fluid, using a BD FACSAria III (BD Biosciences) set to a 70 µm nozzle, 70 PSI sheath pressure, FSC-H 1000, SSC-H 1000, and UV1-H 800. We sorted 6300 nuclei as input for 10x Genomics scDNA library preparation. A total of 6300 nuclei were collected in 4.2 µL 1x PBS (0.1% BSA).

Nuclei for scRNA-seq/scISO-seq library preparation were DAPI-stained (1-5 mg/ml) and sorted in BD FACSFlow™ (BD Biosciences) using a BD FACSAria Fusion Cell Sorter (BD Biosciences) set to a 70 µm nozzle, 70 PSI sheath pressure, FSC 1000, SSC 1000, and UVB 800. A total of 17900 nuclei were collected in 5 µL 1x PBS (0.1% BSA).

**10x Genomics Library Preparation and Sequencing**

scDNA-seq libraries were prepared according to the 10x Genomics user guide Chromium Single Cell DNA Reagent Kits (RevB) and scRNA-seq libraries according to the user guide Chromium Single Cell 3' Reagent Kits v3 (RevC), using the 10x™ Chromium controller. scDNA-seq libraries were sequenced using Illumina HiSeq3000 in the 2x150 bp paired-end mode, whereas scRNA-seq libraries were sequenced using Illumina NovaSeq 6000 in the 2x150 bp paired-end mode.

**Single-cell PacBio ISO-seq**

To create scISO-seq libraries, we used an aliquot of the full-length single-cell cDNA (created with Chromium Single Cell 3' Reagent Kits v3 (RevC) as input from 10x Genomics) for the SMRTbell® Express Template Prep Kit 2.0 from PacBio. This cDNA had the same barcodes as the scRNA-seq. The library was prepared according to the manufacturer's instructions.

#### Whole genome sequencing of leaves

For three branches, a single leaf each was ground with pestle and mortar. The DNA was extracted with Machery Nagel Kit NucleoSpin PlantII. Elution with 40µl PE elution buffer. After the first elution, it was eluted again with the eluate. For the quality of the DNA, the OD260/280 and OD260/280 were measured with NanoDrop. For the exact concentration, the DNA was measured with Qubit. The extracted DNA samples were sent to Novogene for PCR-free library preparation and sequencing using Illumina paired-end sequencing platforms.

#### Somatic mutation validation using digital PCR

We designed ~20 bp primers (fragment size of 80-150 bp) with ~50% GC content and melting temperature ($T_m$) of 58-60°C (Supplementary Table S7). Primers were synthesized and lyophilized by Sigma-Aldrich® and then diluted with nuclease-free water at a 100 µM concentration. For each position, wild-type and mutant probes (length: 20-30 bp) were generated with $T_m$ 5-10°C higher than the primer pair but lower than 72°C. The probes were within 5-10 bp of one of the two primers, non-palindromic, with ~50% GC content, and without G nucleotides at the 5' end. Probes included two locked nucleic acids (LNA), one positioned directly at the SNV site and the other adjacent to the SNV. For two positions CUR6G:15553688 (ID29) and CUR6G:19101663 (ID31), we designed probes for the two wild-type heterozygous alleles and the somatic mutation allele.

Primers were validated using PCR with the SuperFi II Taq Polymerase Green Master Mix protocol (ThermoFisher Scientific). The PCR program consisted of an initial denaturation at 98°C for 30s, followed by 30 cycles of denaturation at 98°C for 10s, annealing at 60°C for 15s, extension at 72°C for 10s and a final extension step at 72°C for 5 min. PCR products were resolved using gel electrophoresis on a 1.5% agarose gel at 100V for 90 min using TBE 1X as the running buffer and visualized using ethidium bromide with BioRad ChemiDoc™ XRS+. Once the fragment amplification was confirmed PCR amplicons were purified using the commercial kit NucleoSpin Gel and PCR cleanup (Macherey-Nagel). Purified PCR products were quantified using

NanoDrop1000 and then sequenced with Sanger technologies by the LightRunXP service of Eurofins Genomics (Germany). Sequences were then analyzed using UGENE software (34).

The digital PCR (dPCR) assays were performed on the QIAcuity platform (*i.e.,* machine and software) using a 26K Nanoplate (24-well) with 26000 partitions. The reaction setup consisted of a final concentration of Qiagen's 1X Probe PCR Master Mix, a primer-probe mix that included 0.8µM forward primer, 0.8µM reverse primer, and 0.4µM probe, 0.25U/µl of PvuII restriction enzyme for genomic DNA, and 20ng of template DNA (approx. 80,000 copies). Thermocycling conditions consisted of an initial heat activation of 2 min at 95°C, 40 cycles of 15s denaturation at 95°C, and a 30s combined annealing/extension at 60°C. This protocol was taken from QIAcuity User Manual Extension: Application Guide.

Results were evaluated by endpoint measurements on the QIAcuty Qiagen machine, by imaging the plate once the cycles finished. Exposure times for imaging steps consisted of 1000ms, 700ms and 500ms for probes imaged on the green, yellow, and crimson probe channels, respectively. The fluorescence levels of excitation and emission of the probes are shown in Supplementary Table S9.

#### Genotyping fruit:L1 and fruit:L2 mutations in leaves

We selected all SNVs identified in the two layers. We did not use indels because genotyping indels at low read counts is highly error-prone and would bias the analysis. Overall, we genotyped 50 and 40 SNVs identified in the L1 from L2 from the fruit, respectively. Of these, three were shared between the layers. Further, as some SNVs were present in multiple branches, we genotyped each SNV in every branch (in which it is identified). This resulted in 99 and 51 genotyping tests for L1 and L2, respectively (Supplementary Table S3). For each read-support cutoff, we counted the number of tests (SNV-branch pairs) in which the SNV is present either only in the layer (L1 or L2) or both the layer and the leaf samples from the branch. We normalised the counts using the number of tests for the L1 and L2, respectively, and the percentage values were plotted.

### Supplementary Figures

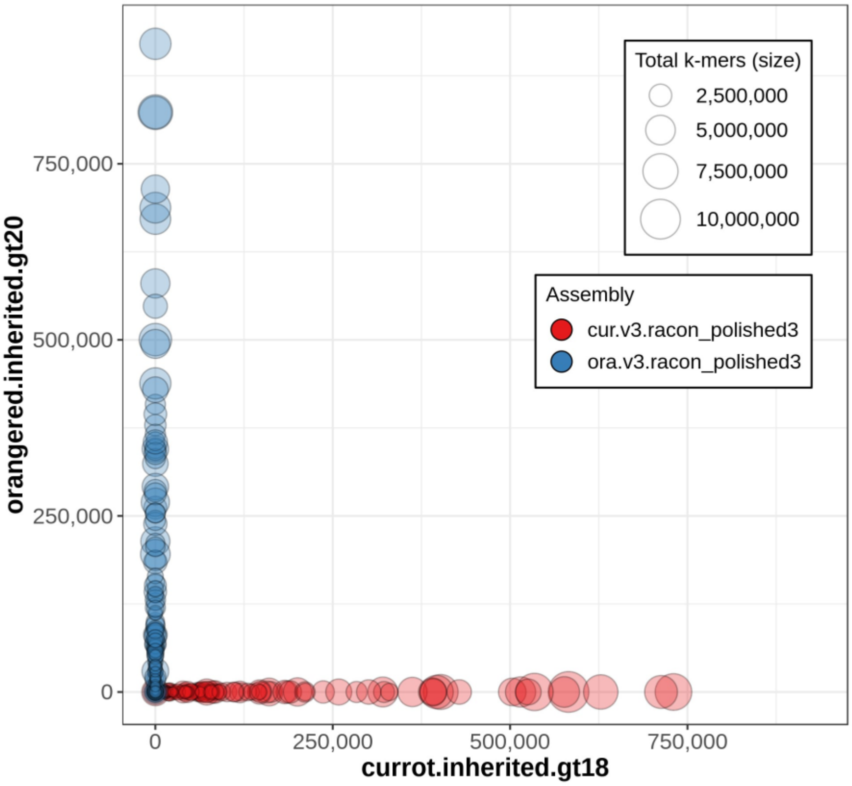


Supplementary Figure S1 *K*-mer-based phasing accuracy of contigs. Contigs from haplotype 1 (Currot) and 2 (Orange Red) are in red and blue respectively. Haplotype-specific k-mers map only to contigs from the matching haplotype implying high phasing accuracy. Plot generated using Merqury.

Currot linkage groups

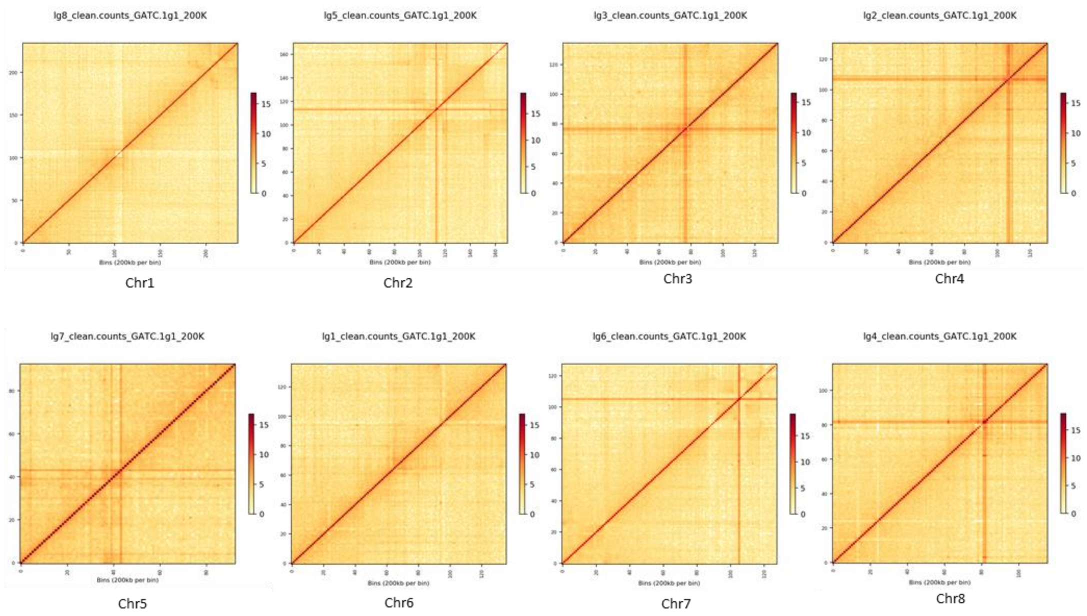


Supplementary Figure S2 Hi-C linkage for chromosomes of haplotype 1 (Currot).

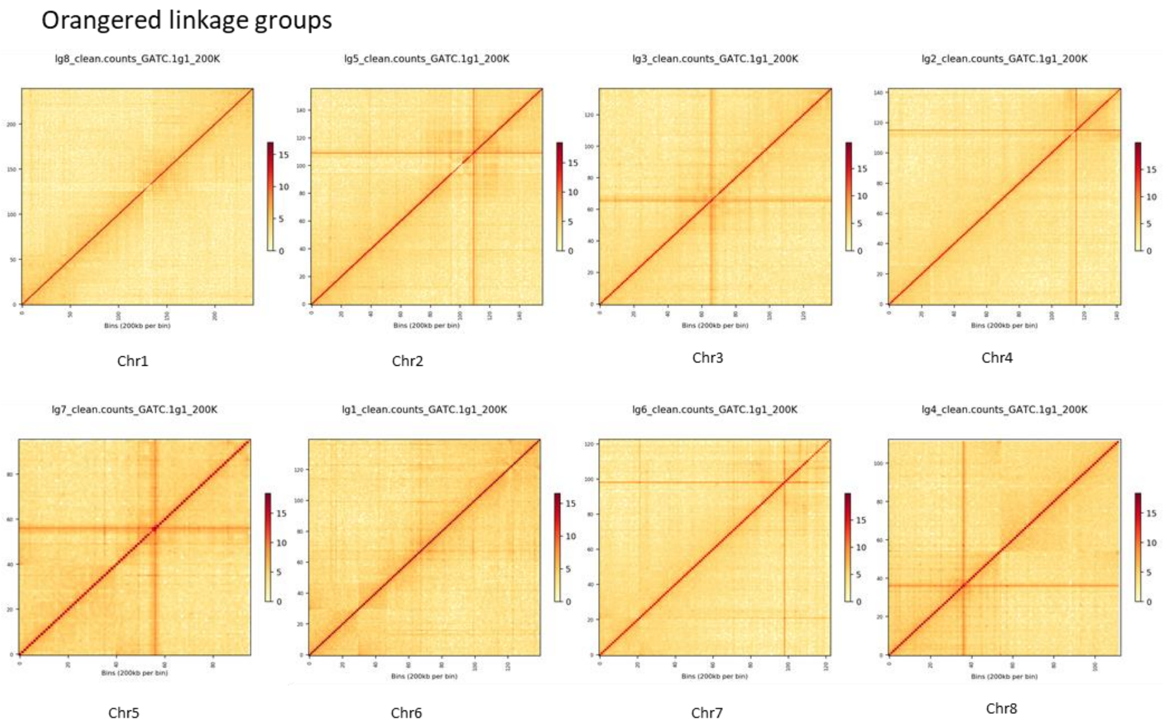


Supplementary Figure S3 Hi-C linkage for chromosomes of haplotype 2 (Orange Red)

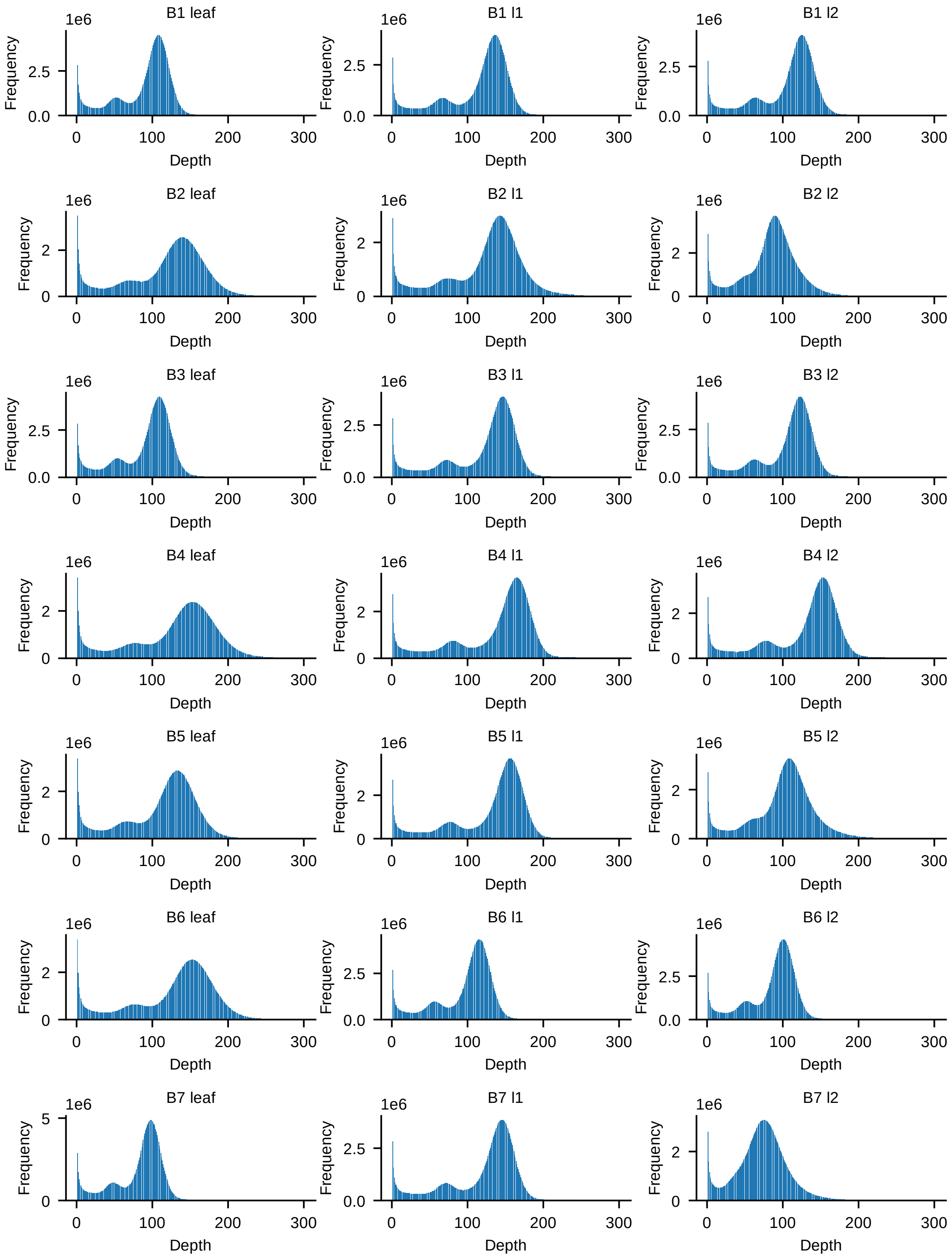


Supplementary Figure S4 Read depth distribution after filtering based on mapping quality and base quality.

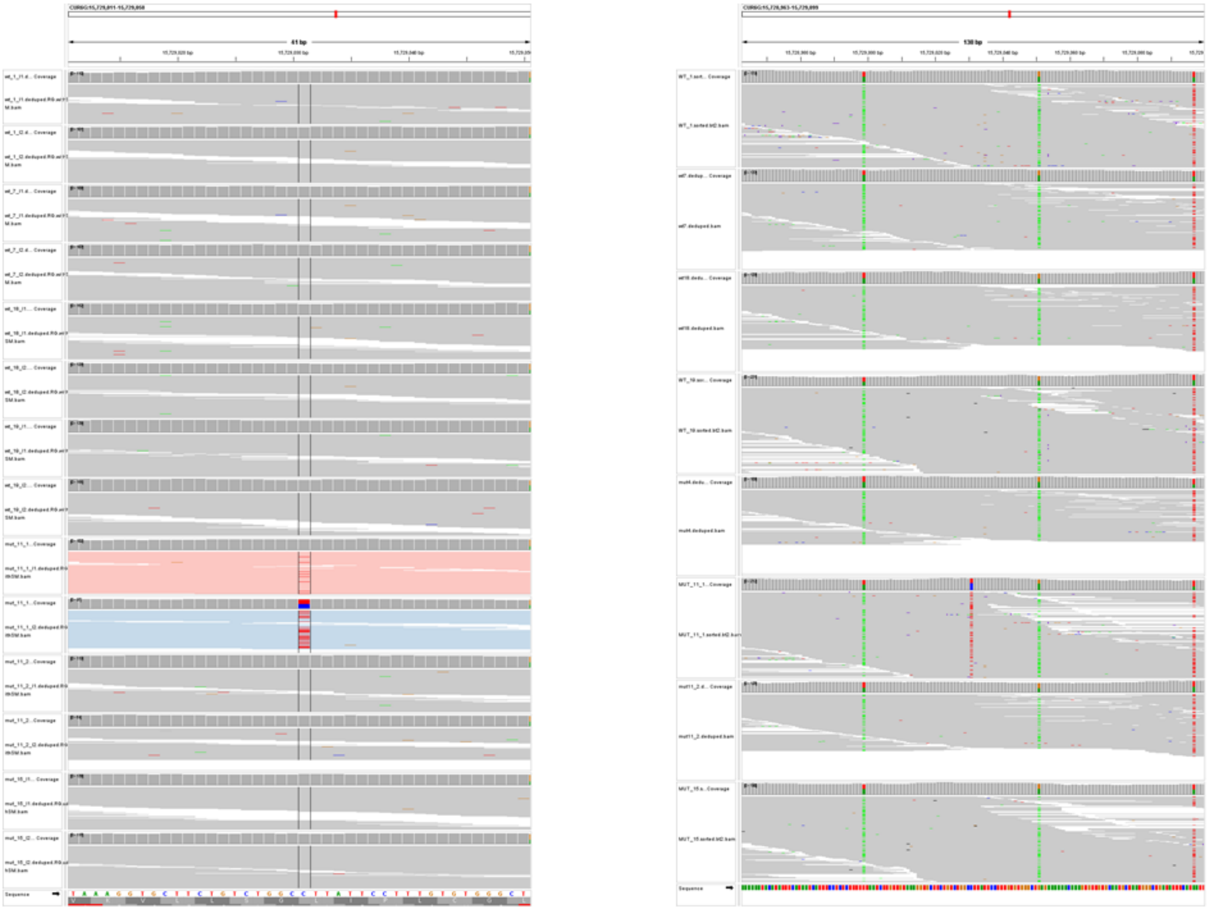

Supplementary Figure S5 Example of somatic mutations. The left panel shows IGV read alignments for 14 layer-sequencing datasets from fruits. Reads from branch B6 are coloured in pink (L1) and blue (L2). The somatic mutation is present only in B6:L2 with some spill-over reads in B6:L1. The right panel has seven leaf samples with a somatic mutation in only one sample.

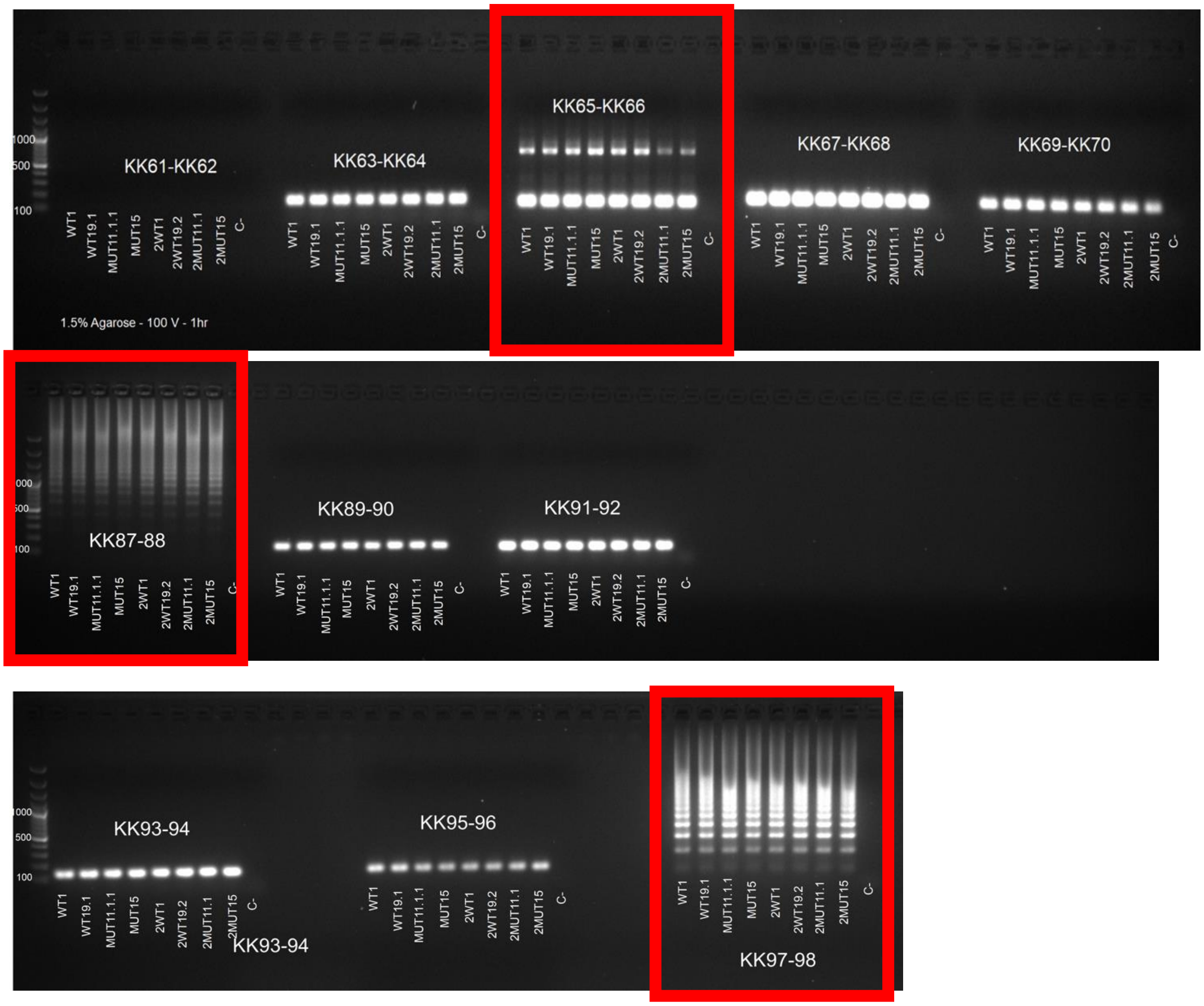


Supplementary Figure S6 Gel electrophoresis images for some somatic mutations tested with digital PCR. The three red boxes highlight positions where the primers did not work.

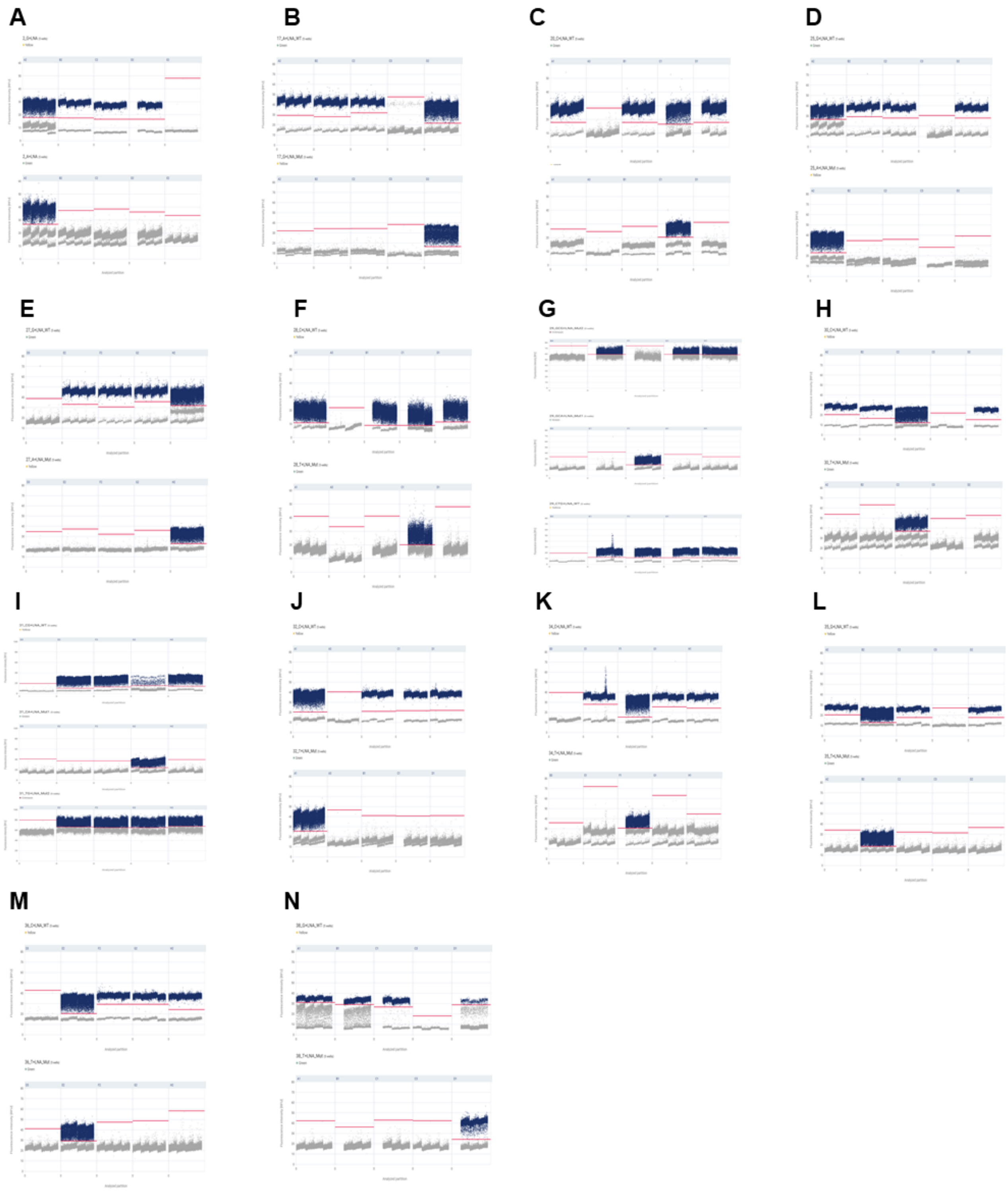


Supplementary Figure S7 dPCR results for the 14 validated somatic mutations. For each subplot, the first (upper) scatterplot is for the wild-type probe and the second (lower) is for the mutant probe. The third scatterplot is for somatic mutations that have a linked heterogenous position. A) ID2, B) ID17, C) ID20, D) 25, E) ID27, F) ID28, G) ID29, H) ID30, I) ID31, J) ID32, K) ID34, L) ID35, M) ID36, and N) ID38. Supplementary Table S10 lists the chromosome positions for dPCR IDs.

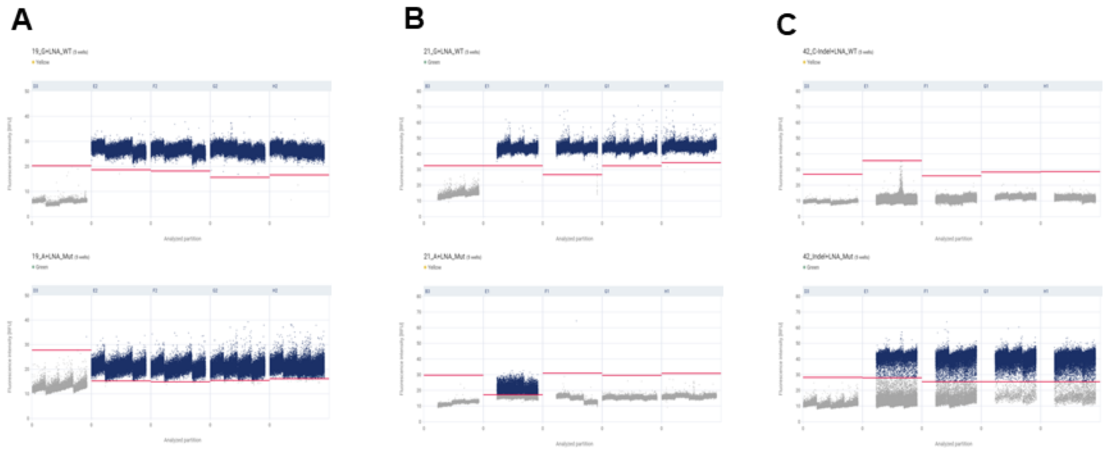


Supplementary Figure S8 Scatterplots displaying non-conclusive patterns on mutations **A.** ID19: no positive/negative partition distinction **B.** ID21: no positive/negative partition distinction **C.** ID42: partition smear. Samples from left to right: Negative control, B2, B4, B5, and B6.

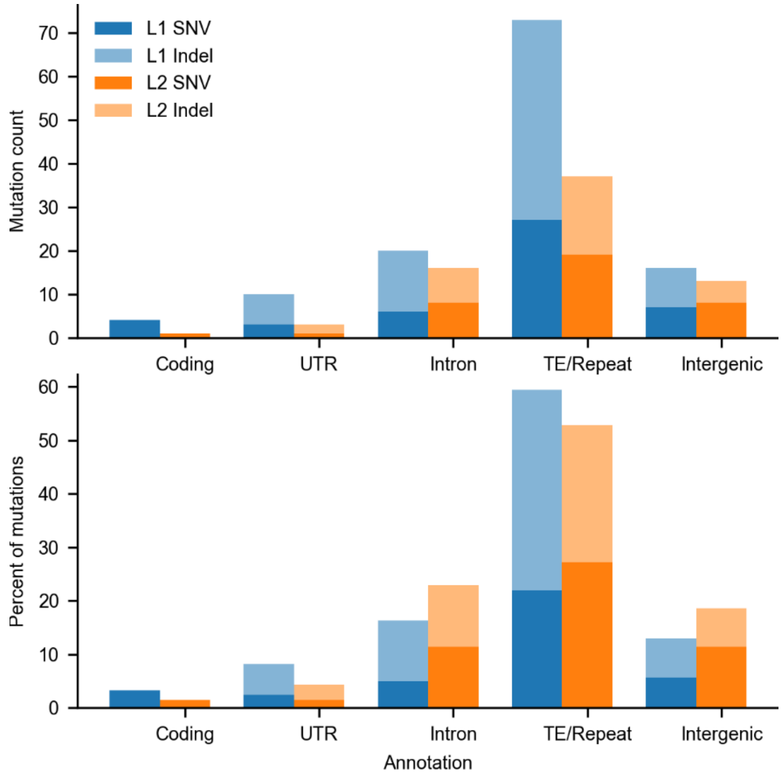


Supplementary Figure S9 Somatic mutations in different genomic regions.

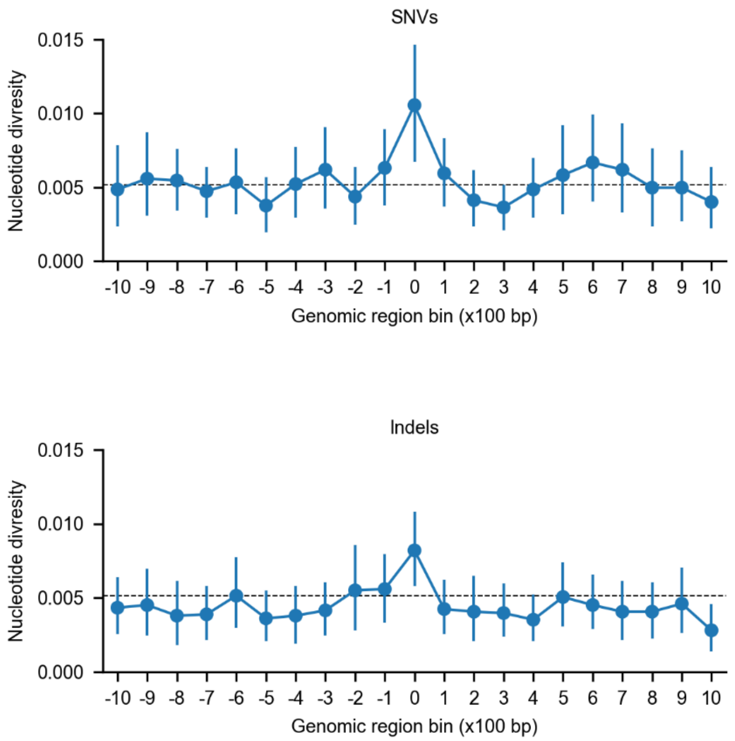


Supplementary Figure S10 Nucleotide diversity upstream and downstream of the identified somatic mutations (SNVs and indels). The dashed line corresponds to average nucleotide diversity. The points are mean nucleotide diversity and the error-bars show confidence interval.

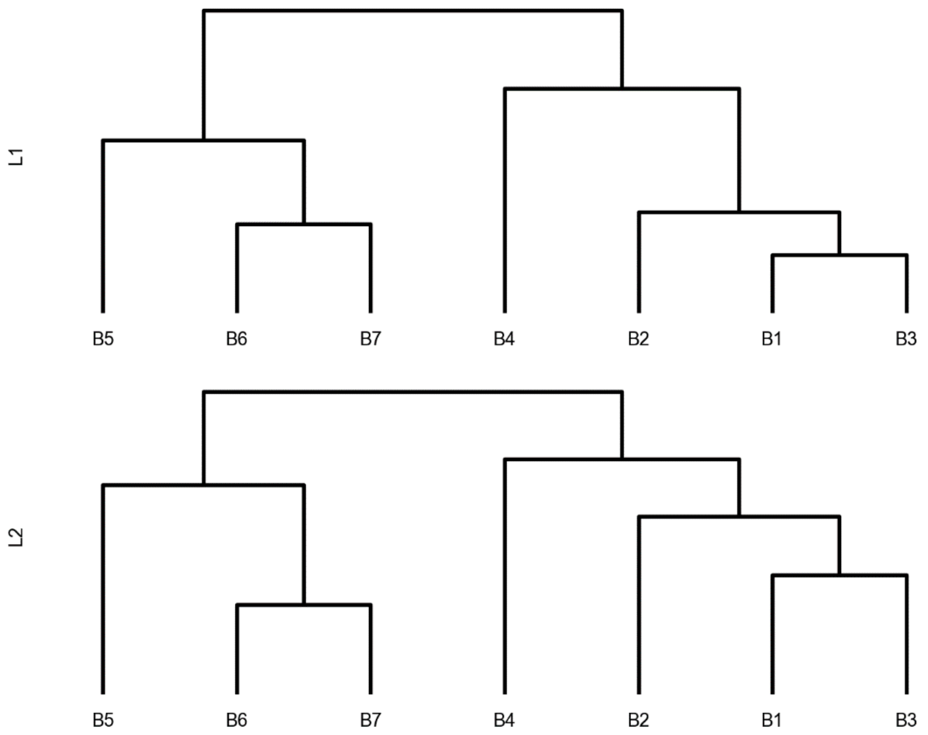


Supplementary Figure S11 Hierarchical clustering of branches based on somatic mutations present in L1 (upper panel) and L2 (lower panel). This matches the tree topology with the exception of B1-B3 clustering together instead of B1 clustering with B2. This is caused by a high number of somatic mutations in B2.

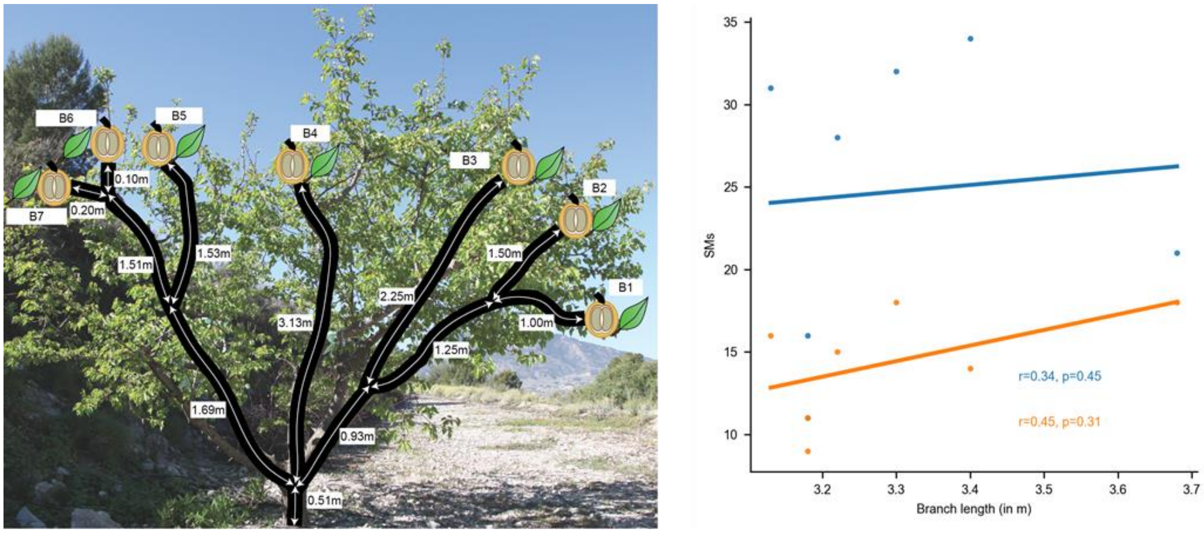


Supplementary Figure S12 Correlation between branch lengths and somatic mutations. The left panel shows branch length of individual branches. The right panel shows the correlation for L1 (blue) and L2 (orange) mutations.

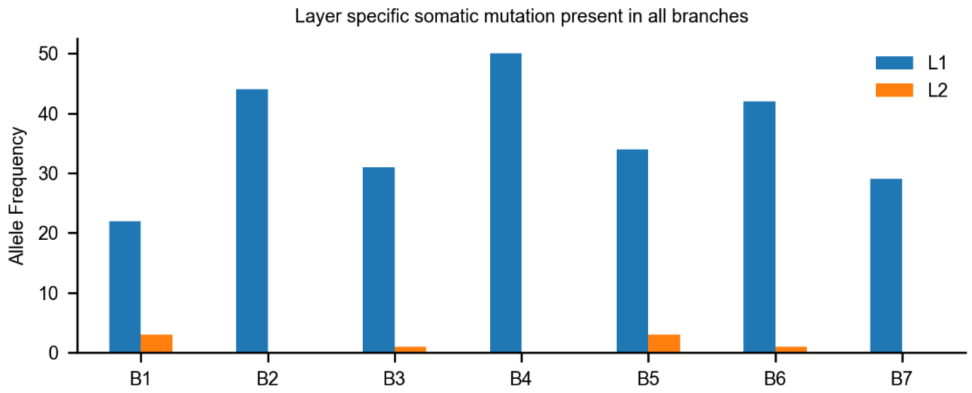


Supplementary Figure S13 Allele frequencies for a somatic mutation present in L1 of fruits from all branches but absent in L2.

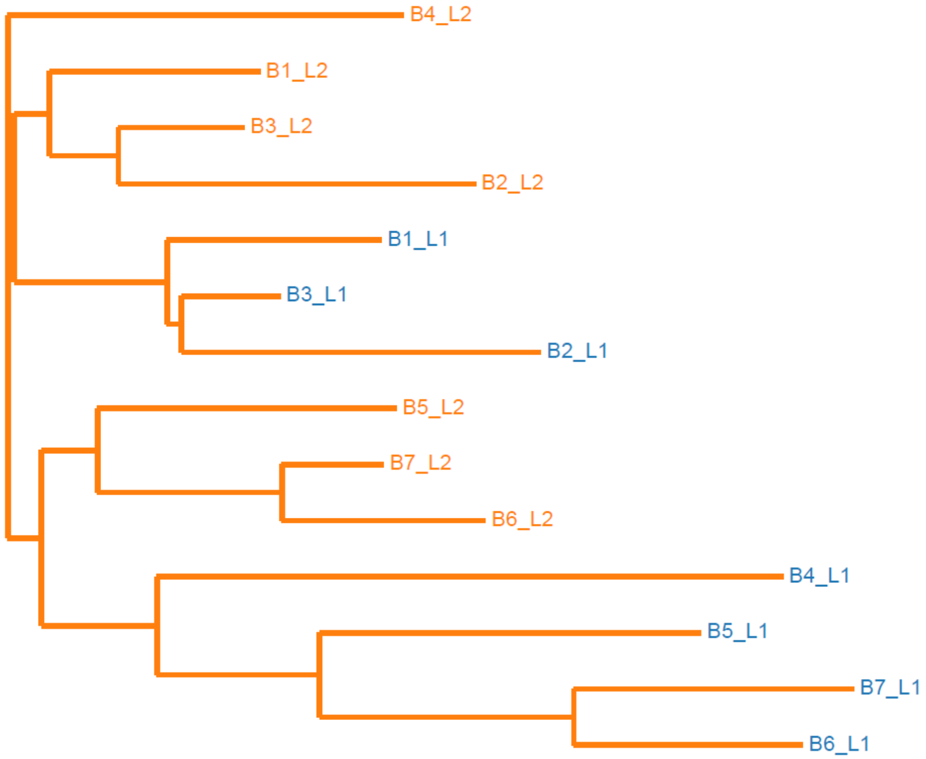


Supplementary Figure S14 Phylogenetic relationship between all samples after removing putatively clonally inherited mutations present in all branches. Somatic mutations occurring in branches formed after the primary branching event cannot be transmitted to other branches. Consequently, samples from the subtree are more similar to each other.

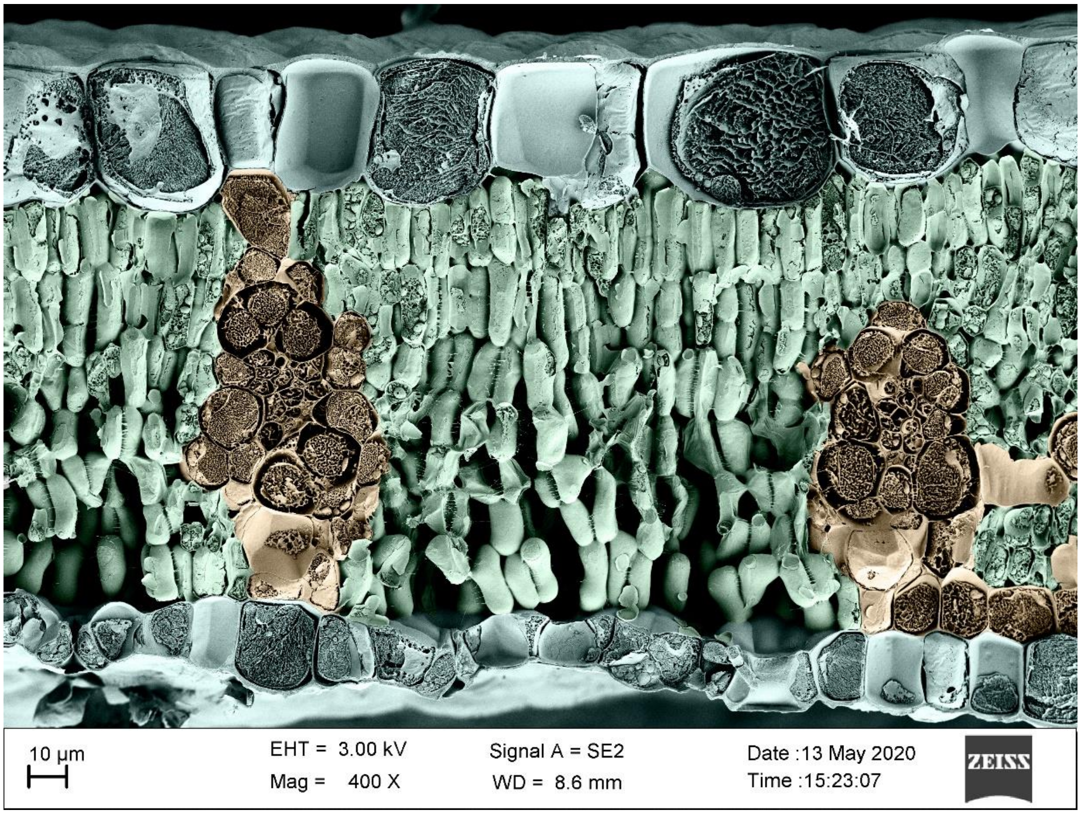


Supplementary Figure S15 Cross-section of an apricot leaf. Grey cells are epidermis (L1), green cells are mesophyll (L2), and brown cells are vascular (L3).

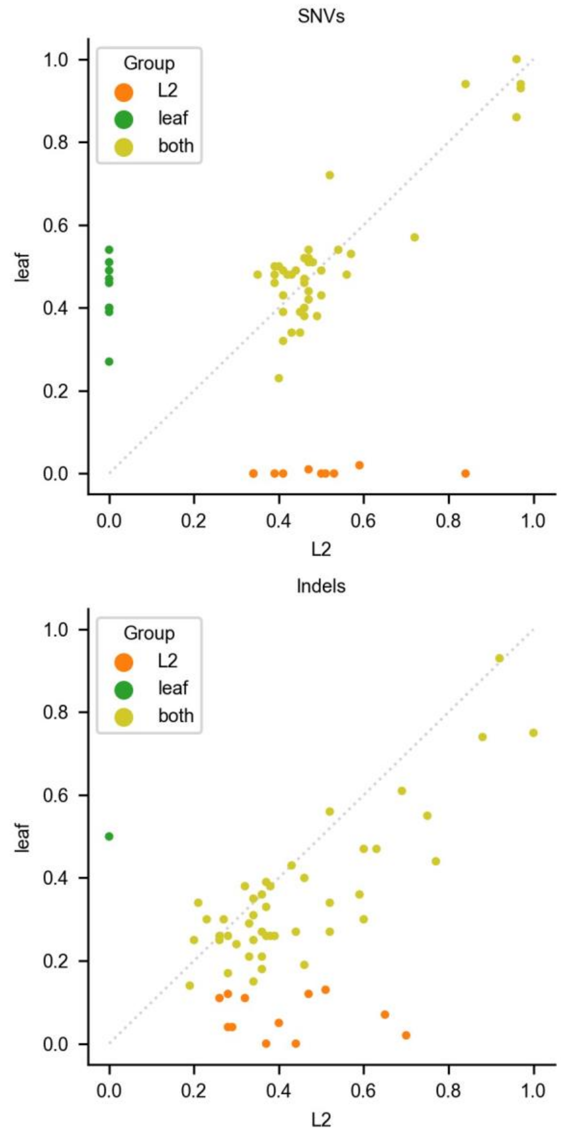


Supplementary Figure S16 Allele frequencies of somatic mutations identified in only L2 (orange), only leaves (green), and in both (golden). X-axis has allele frequency for Fruit:L2 and y-axis has allele frequency for leaf.

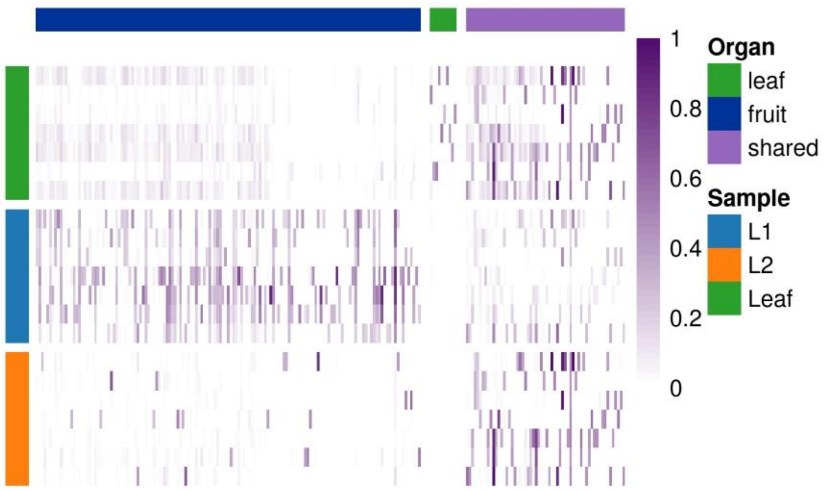


Supplementary Figure S17 Heatmap for allele-frequencies at the identified somatic mutation positions. The rows correspond to samples grouped by leaves (green), L1 from (blue), and L2 from fruit (orange). The columns correspond to SMs grouped based on the organs in which they are called. SMs only in only fruit have violet annotation, only in leaf have green, and shared in both have purple. Alelle-frequency (pruple spectrum) ranges between 0 and 1.

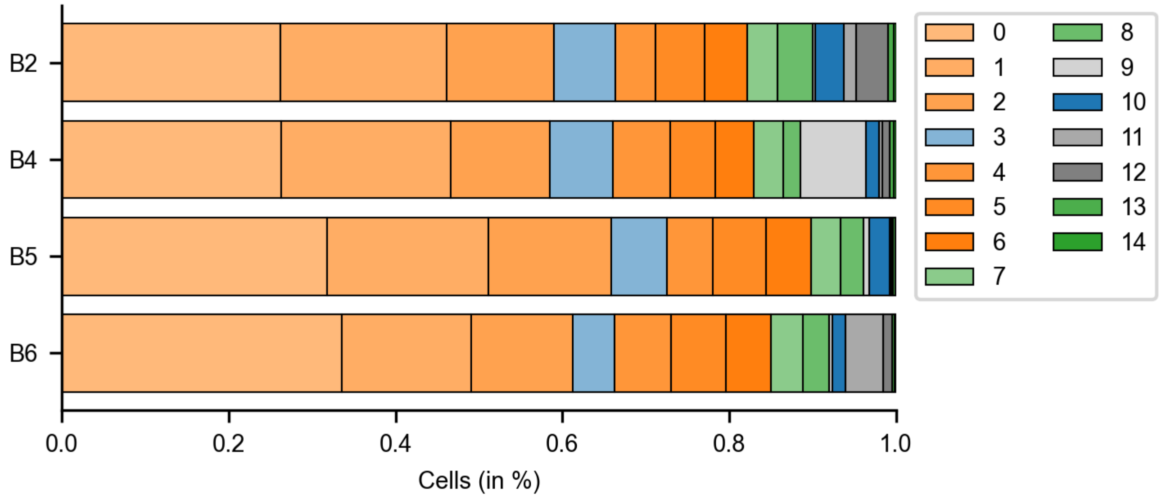


Supplementary Figure S18 Distribution of cells in different clusters. Each partition corresponds to a cell cluster. Clusters corresponding to: L1 are in shades of blue, L2 are in shades of orange, and L3 are in shades of green. Cells corresponding to stress response and dividing cells are in shades of grey.

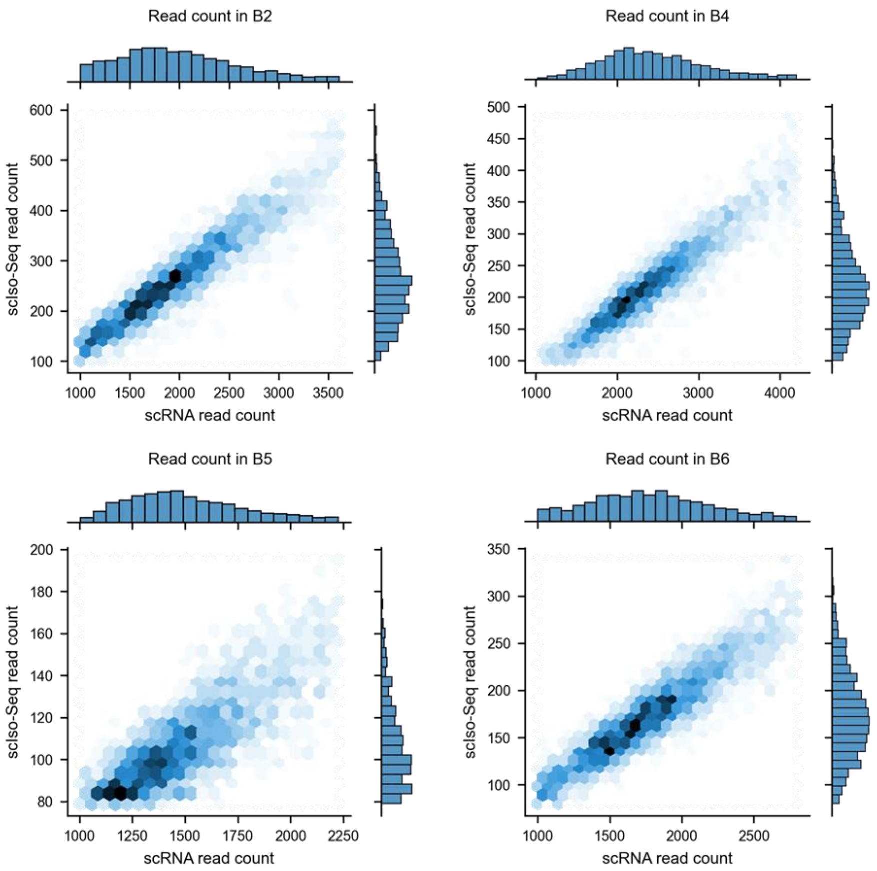


Supplementary Figure S19 Correlation between read-counts in the scRNA-seq and scISO-seq datasets. Reads were paired based on matching barcodes.

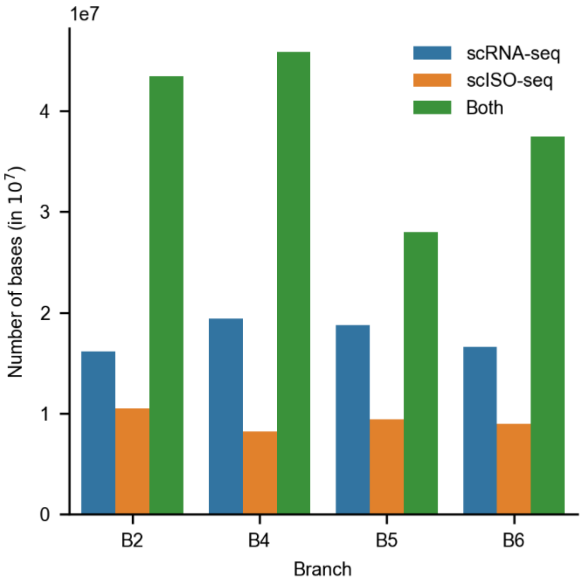


Supplementary Figure S20 Number of bases sequenced by the two technologies

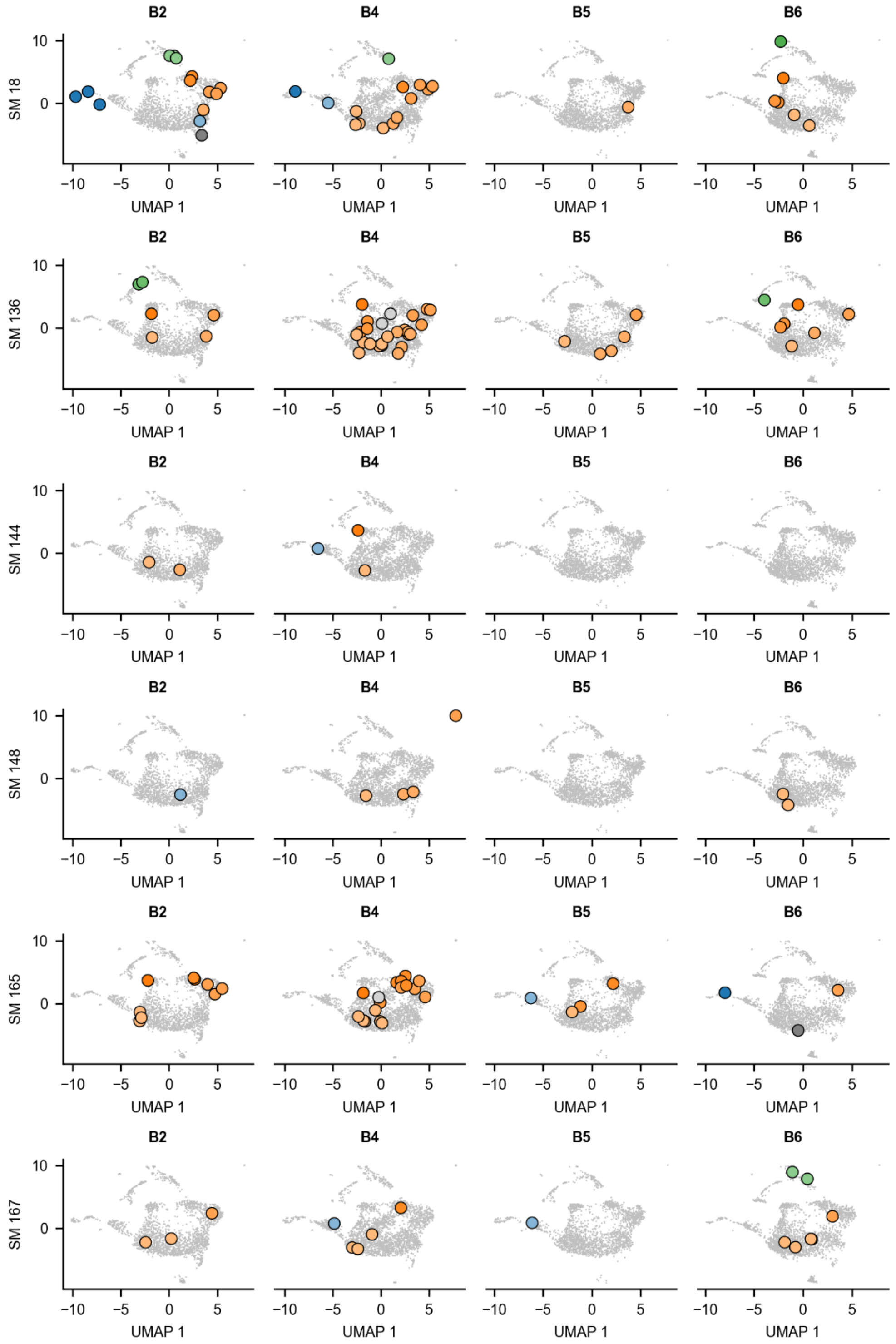


Supplementary Figure S21 Cells containing wild-type alleles for transcribed somatic mutations. The four columns correspond to four branches. Each row corresponds to one somatic mutation. Grey cells do not have any read with the wild type allele. Remaining cells are colours based on the cluster in which they are from.

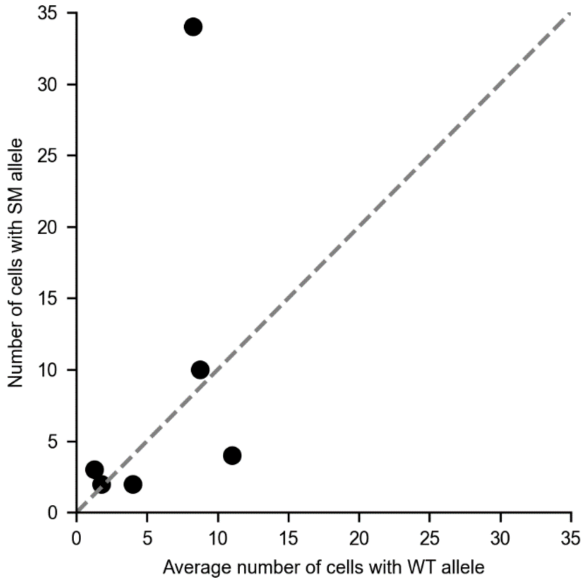


Supplementary Figure S22 The number of cells with mutant alleles were proportional to number of cells with wild-type allele. Each point corresponds to one somatic mutation.

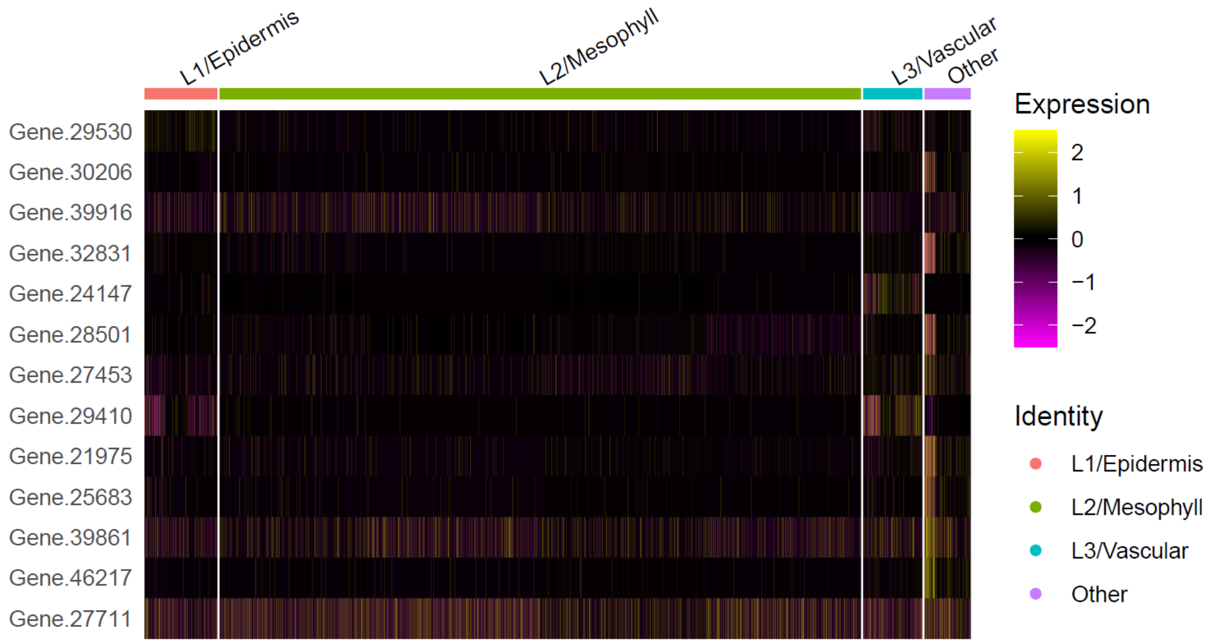


Supplementary Figure S23 Normalised expression values for DNA repair genes in scRNA-seq data. Only *FindMarkers* genes with adjusted p-values less than 0.05 are shown.

### Supplementary Tables in Additional File 2


| | |
|---|---|
| Table S1 | Raw whole-genome sequencing reads summary |
| Table S2 | Read depth cutoffs used for variant calling |
| Table S3 | List of identified somatic mutations |
| Table S4 | Allele frequency variation at SM position in all whole-genome sequencing samples |
| Table S5 | List of gene conversions |
| Table S6 | List of identified complex variations |
| Table S7 | Pair of primers used for dPCR |
| Table S8 | Pair of probes designed for each candidate mutation |
| Table S9 | Channels used on dPCR assays and its exposure times during imaging step |
| Table S10 | Somatic mutations validated using dPCR |
| Table S11 | Manually curated list of marker genes |
| Table S12 | Annotation of single cell clusters |
| Table S13 | All gene markers identified from the scRNA-seq cluster analysis |
| Table S14 | List of gene markers with layer-specific functional annotation |
| Table S15 | Enrichment analysis for cell clusters |
